# Governing the decline: clam fisheries and the challenges of decentralized management across the western Mediterranean and Gulf of Cádiz (Spain)

**DOI:** 10.64898/2025.12.03.692034

**Authors:** Marc Baeta, Laura Benestan, Mauricio Mardones, Marina Delgado, Luis Silva, Miguel Rodilla, Silvia Falco, Manuel Ballesteros, Miriam Hampel, Ciro Rico

## Abstract

Over the past four decades, clam fisheries along the Mediterranean and Atlantic coasts of Spain have exhibited significantly different ecological and governance trajectories. This study synthesises long-term landing data from 1993 to 2024 with a systematic review of regional legislation to explore how management frameworks influence the resilience of fisheries. In the northwest Mediterranean, which includes Catalonia, the Valencian Community, the Balearic Islands, and Murcia, fisheries have experienced synchronous declines, a consequence of overexploitation, fragmented governance, and reactive, top-down regulation. In contrast, Andalusia, encompassing both the Atlantic and Alboran Sea coasts, has demonstrated more resilient trajectories, bolstered by greater ecological productivity and the adoption of adaptive governance strategies since the 2010s. In this region, the implementation of gear-specific management plans, rigorous scientific monitoring (including satellite tracking), and prompt administrative responses has facilitated more sustainable exploitation. Our findings highlight the necessity of aligning governance with ecological connectivity and integrating adaptive co-governance structures that engage fishers to safeguard the future of small-scale fisheries. More broadly, this research emphasises the need for coordinated superregional or national management frameworks that incorporate ecological knowledge, promote stakeholder participation, and allow for timely regulatory adaptation.

## 1. Introduction

Marine fisheries are a key source of high-quality protein and play a central role in global food security, employment, and trade [1]. Since the mid-20th century, global demand for seafood has grown steadily, driven by population growth, rising incomes, and increasing awareness of its health benefits[2]. However, wild marine fishery production has plateaued at around 90 million tonnes annually since the late 1980s [1]. Aquaculture has expanded rapidly to bridge the widening supply-demand gap, but in the European Union (EU), this imbalance is particularly acute. Over the past three decades, wild fisheries production has steadily declined while seafood demand has continued to increase. Aquaculture output has remained relatively stable and insufficient to offset this decline [1]. Consequently, the EU has become increasingly reliant on seafood imports [4]

The decline in EU fisheries productivity has been particularly pronounced in small-scale fisheries (SSF) [5]. One example is the marked contraction of SSF targeting commercial clams (e.g., *Donax trunculus*, *Chamelea gallina*, and *Callista chione*) along Spain’s Mediterranean and southwestern Atlantic coasts, especially in Andalusia. Once an important source of regional economic activity, supported by high clam landings during the 1990s, these fisheries now persist only in Andalusia and Catalonia (Fig. 1). Multiple factors have been proposed to explain these declines, including environmental variability, fishing pressure, and other anthropogenic pressures [6–8]However, the extent to which governance and management have amplified or mitigated these dynamics remains largely unexplored. The SSF sector still accounts for most of the Spanish fleet, representing 64% of landings between 2006 and 2020. SSF typically involves vessels under 12 metres in length operating in coastal and adjacent waters, crewed by small teams and using diverse fishing gears on daily trips. The sector also includes shore-based shellfishers who target sedentary and low-mobility species, such as clams, cockles, barnacles, and polychaetes, collected during low tide or in shallow waters less than 1.5 metres deep [9–12]. These fisheries often require low initial investment and are generally owner-operated [13]. Compared with industrial and semi-industrial fisheries, SSF show lower costs, catches, turnover, reliance on subsidies, and fuel consumption [14]. Despite their smaller scale, SSFs generate higher profits per unit of investment, greater catch efficiency per litre of fuel, and higher socioeconomic value per kilogram of harvested marine resources [15]. They also play a vital role in employment, local development, the equitable distribution of economic benefits, and the supply of fresh, high-quality seafood to local, regional, national, and international markets [16]. Within this broader context, the persistence of SSF is crucial for employment and for the economy of coastal communities across the EU [17].

**Figure 1.**
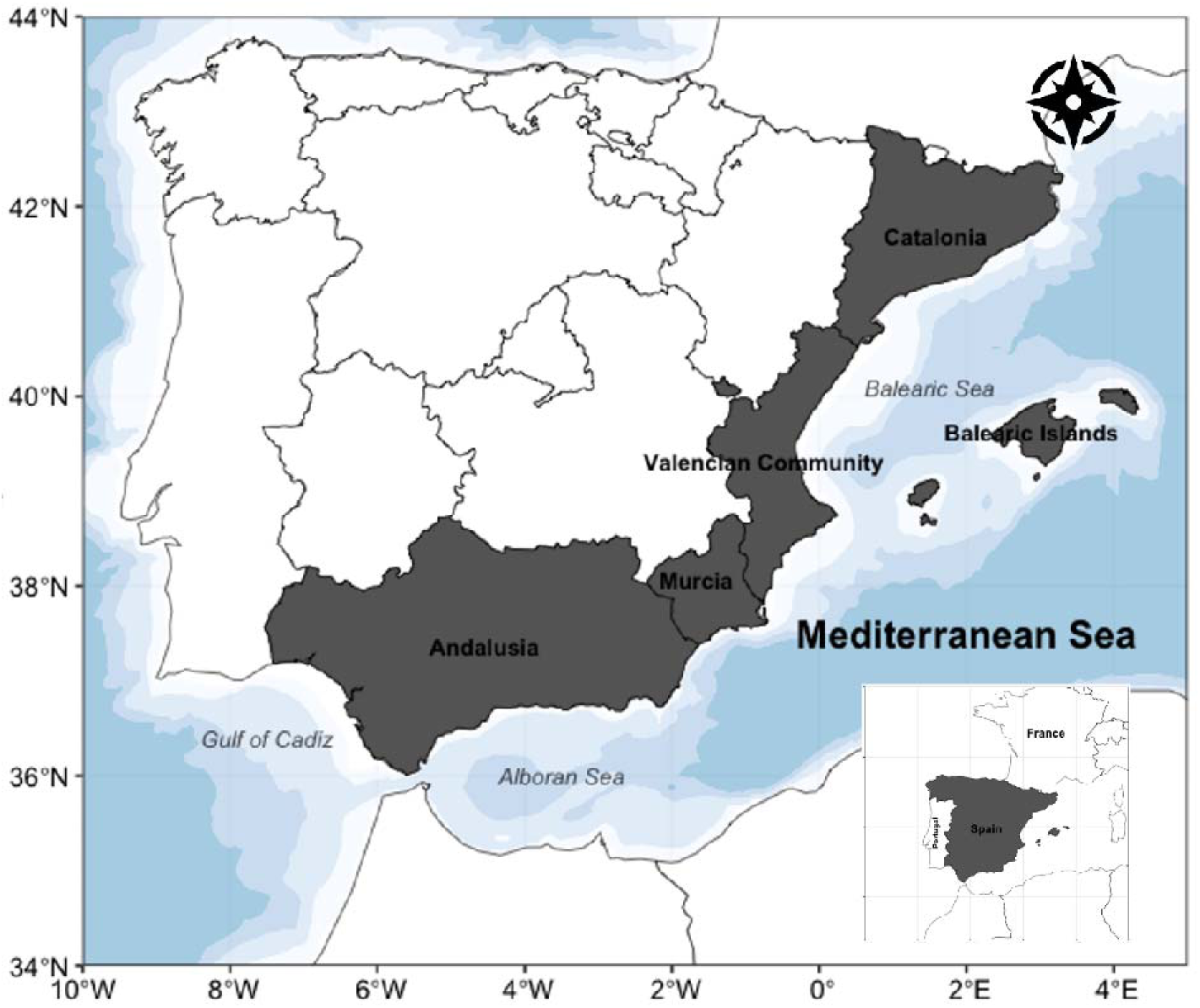
Map of the study area, with the Autonomous Community where the main clam fisheries are located.

Clam species commercially exploited in Spain are shallow-burrowing, suspension-feeding invertebrates that inhabit clean sandy substrates in coastal waters and share a broadly similar geographic distribution. Like many other bivalves, they are sedentary or even sessile and typically show an aggregated distribution [18,19]. Their populations are structured as metapopulations composed of spatially discrete subpopulations connected through pelagic larval dispersal [20]. Wedge clams (*D. trunculus*) inhabit dynamic sandy shorelines [21] and are predominantly found in well-sorted fine sand biocenoses at depths ranging from 0 to 3 m in the Mediterranean Sea [22] and from 0 to 6 m in the Atlantic Ocean [21]. Striped venus clams (*C. gallina*) occupy a broader range of sediment types (sand, sandy mud, and mud), with a preferential distribution in well-sorted fine coastal sands at depths of 3 to 20 m [22], whereas smooth clams (*C. chione*) are found in coarse sands at depths of 5 to 30 m [7,22]. These habitat preferences strongly influence exposure to hydrodynamic disturbance, sediment instability, granulometry, and environmental variability (*e.g.*, temperature and salinity), all of which affect survival, growth, spawning, and recruitment success [7,18,23]. For example, fluctuations in the abundance of *C. gallina* in the Alboran Sea have been linked to environmental conditions, particularly during spawning and recruitment periods [24]. In addition, mass mortality events in adult commercial clams and cockles have been documented in association with disease outbreaks, hypoxia, extreme temperature anomalies, and sediment disturbance [23,25–27]. Such ecological dynamics can produce abrupt fluctuations in landings that are not solely attributable to fishing pressure. Because many commercially exploited sandy-beach clams are relatively short-lived and mature rapidly (e.g., *D. trunculus* and *C. gallina*), fluctuations in recruitment success or mass mortality can translate into pronounced short-term changes in stock size. In such environmentally dynamic and spatially connected systems, the scale and design of governance frameworks may critically influence the capacity to respond effectively to ecological fluctuations. This potential ecological-institutional mismatch constitutes the central analytical lens of the present study [28,29].

SSF vessels and shore-based shellfishers targeting clams generally operate in nearshore waters. Since the 1980s, coastal fisheries management in Spain has been decentralised [30], giving autonomous communities (Comunidades Autónomas, AACC) full authority over regulatory and licensing frameworks. These frameworks govern fishing permits, designated fishing areas, quotas, allowable fishing days, and operating schedules. However, despite the diversity of governance models and regulatory systems adopted by each AACC, most clam fisheries have either collapsed or are now experiencing marked declines [6,18,31–33]. Fisheries governance encompasses both formal and informal regulations, institutions, and decision-making processes that define access to and exclusion from marine resources and shape how stakeholders influence their use and management [34]. The distribution of commercial clam species is often patchy, with dense populations restricted to sites where environmental conditions are favourable. As a result, fishing effort tends to concentrate in relatively small areas. Yet the natural biogeographical boundaries of clam populations frequently do not align with administrative borders. For example, Marie, [35] identified three distinct populations of *D. trunculus*: one spanning the Catalan and Valencian AACCs, another along the Mediterranean coast of Andalusia, and a third along the Atlantic coast of Andalusia. This implies that different regulatory regimes and fishing pressures have historically affected a single Northwestern Mediterranean population across its range. To date, research on SSF targeting clams has remained largely local, focusing on individual AACCs in Catalonia, the Valencian Community, and Andalusia [18,31,36–38]. No study has yet examined these fisheries at a broader regional scale while explicitly considering the interaction between ecological variability and decentralised governance systems. Given the environmental sensitivity of these species and the fragmentation of management across AACCs, a regional perspective is essential for understanding why some fisheries have persisted while others have collapsed. This study therefore provides the first integrated regional analysis of commercial clam fisheries along the Spanish Mediterranean and southwestern Atlantic coasts, combining long-term landing data with a comparative assessment of governance structures to examine how ecological dynamics and institutional design have jointly shaped divergent trajectories.

## 2. Material and Methods

### 2.1 Study area

Spain is politically divided into regions known as Autonomous Communities (AACC) that enjoy significant legislative, executive, and administrative power [40]. Along the Mediterranean coast, these include Catalonia, the Valencian Community, the Balearic Islands, Murcia, and Andalusia (Fig. 1). This 2,200 km coastline contains diverse ecosystems shaped by both natural processes and human activities. It is subject to substantial anthropogenic pressure, particularly because of high population density and seasonal tourism during the summer months [39]. Industrial centres, including Barcelona, Tarragona, Castellon, Valencia, Alicante, Cartagena, Almeria, and Malaga, contribute to localised pollution, whereas agricultural runoff has a more diffuse but comparatively limited impact [40].

In contrast, the Atlantic coast of Andalusia represents a distinct ecological and socio-political region, extending approximately 300 km from the Strait of Gibraltar to the Portuguese border. This area is strongly influenced by Atlantic oceanographic conditions, including higher productivity, stronger tidal regimes, and seasonal nutrient inputs from major rivers such as the Guadalquivir and Guadiana [41,42]. It is also subject to considerable human pressure, particularly from summer tourism, rapid coastal urbanisation, and intensive agriculture in the lower Guadalquivir basin and around Doñana National Park [43]. Moderate to high environmental risks from pesticide contamination have been documented in this area [44] Despite these pressures, the most pronounced declines in clam landings have been observed in the Northwestern Mediterranean Sea, which is therefore the primary focus of this study.

### 2.2 Data collection and analysis

This study combined official landing statistics and regulatory documentation to assess long-term trends and regional governance approaches in Spanish clam fisheries. Annual landings (1993–2024) for the three target species (*D. trunculus*, *C. gallina*, and *C. chione*) were obtained from the regional Fisheries Directorates of the AACCs (Catalonia, the Valencian Community, the Balearic Islands, the Region of Murcia, and Andalusia). These datasets were supplemented with information from statistical yearbooks, grey literature, and peer-reviewed publications to ensure temporal continuity and reliability. To evaluate regional differences in management, we conducted a systematic legal and documentary review. For each AACC, we compiled and analysed all available legal texts related to the management of the target species, including regional decrees, ministerial orders, management plans, and administrative resolutions issued between the early 1980s and the 2020s. Regulatory documents were identified through searches of official regional government bulletins and fisheries administration websites, complemented by grey literature and institutional reports. The documents were analysed qualitatively to identify the timing, scope, and type of governance measures implemented in each region, including licensing systems, effort controls, spatial regulations, seasonal closures, monitoring schemes, and management plans.

Non-parametric statistical tests were used to examine temporal patterns and relationships in clam landings across Spanish coastal regions from 1993 to 2024. Spearman’s rank-order correlation coefficients were calculated to evaluate: (1) the relationship between total annual landings by species (*C. gallina*, *D. trunculus*, and *C. chione*) and the Autonomous Communities (AACC), including Catalonia, the Valencian Community, the Balearic Islands, Murcia, Mediterranean Andalusia (Alboran Sea), and Atlantic Andalusia (Gulf of Cadiz); and (2) the relationships among species’ landings across three geographic areas: the Northwestern Mediterranean Sea, Mediterranean Andalusia, and Atlantic Andalusia. For the statistical analysis, the Northwestern Mediterranean region includes Catalonia, the Valencian Community, the Balearic Islands, and the Region of Murcia. This aggregation was used only to explore possible synchronies in landing trends within the basin and does not imply similar governance frameworks across these regions, which are examined separately in the governance analysis. These analyses were intended to identify synchronies in temporal trends, regional disparities in exploitation patterns, and economic signals related to commercial demand. Correlations were considered statistically significant at a 99.9% confidence level (P < 0.001).

Environmental variability was examined to assess potential relationships between environmental conditions and annual landings of the three species of clams analysed in this study, namely *D. trunculus*, *C. gallina*, and *C. chione*. Environmental data were obtained from the Copernicus Marine Service (CMEMS) database for the period 1993–2024. Three environmental variables were considered: sea surface temperature (SST), sea surface salinity (SSS), and chlorophyll-a concentration (CHL). SST and SSS were derived from CMEMS ocean reanalysis products, whereas chlorophyll-a data was obtained from satellite-derived ocean-colour products. Environmental data were extracted for three regional domains corresponding to the main fishing areas analysed in this study: the NW Mediterranean (43.0°N–38.5°N, 0.5°W–4.5°E), the Alboran Sea (37.5°N–35.5°N, 5.5°W–1.5°W), and the Gulf of Cádiz (37.5°N–35.5°N, 9.5°W–5.5°W). Monthly data were extracted for each region and aggregated to annual means. Environmental anomalies were calculated as deviations from the long-term mean for the study period. To explore potential recruitment effects, SST and SSS were analysed using a one-year lag relative to annual landings. Pearson correlation analyses were then performed between annual landings and environmental variables (SST_lag1, SSS_lag1, and CHL anomalies) for each species-region combination. Environmental anomaly time series are shown in the Supplementary Material (Fig. S1), and correlation coefficients and associated p-values are provided in Table S1.

To integrate the environmental and governance dimensions of the study, we applied a comparative socio-ecological framework. Long-term landing trajectories were interpreted in relation to the management measures and governance frameworks documented for each AACC. This comparative approach allowed us to identify regional patterns in the interaction between fishery dynamics, environmental variability, and governance responses. Although it does not establish direct causal relationships, it provides an integrative perspective on how environmental variability and institutional responses may jointly shape long-term fishery trajectories.

## 3. Results

### 3.1. Landings

The findings revealed strong and consistent positive Spearman correlations within the Northwestern Mediterranean (NW Mediterranean, hereafter) particularly concerning *C. gallina* (see Table 1). This species showed significant correlations across all four AACC (Catalonia, the Valencian Community, the Balearic Islands, and Murcia). Additionally, *D. trunculus* demonstrated a significant correlation between Catalonia and the Valencian Community. In contrast, the correlations between the NW Mediterranean and the southern regions (Mediterranean Andalusia and Atlantic Andalusia) were weak, inconsistent, or non-existent, indicating that landings in these areas follow distinct trajectories. Consequently, landings were categorised and analysed by management area: (1) the NW Mediterranean (comprising Catalonia, the Valencian Community, Murcia, and the Balearic Islands), (2) Mediterranean Andalusia, and (3) Atlantic Andalusia.

**Table 1.**
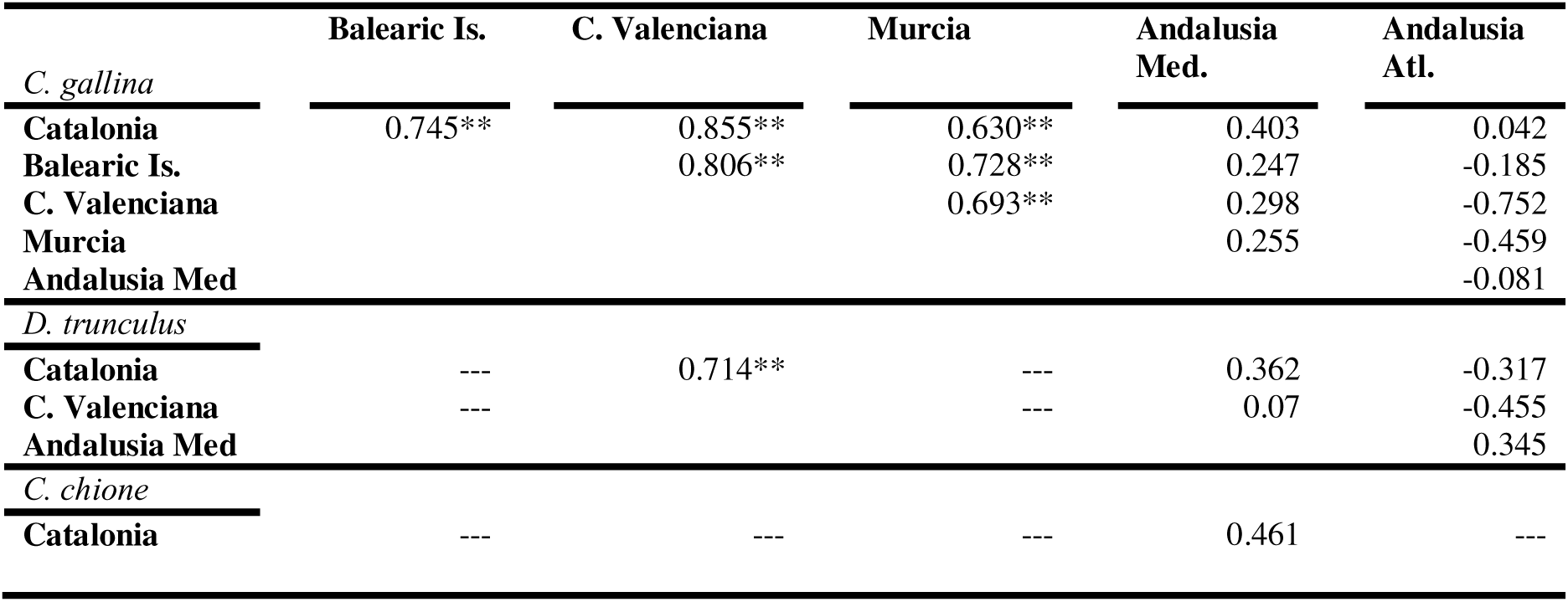
Spearman rank-order correlation coefficients between annual landings of *Chamelea gallina, Donax trunculus*, and *Callista chione* across AACC (Catalonia, Balearic Islands, Comunitat Valenciana, Murcia, Andalusia Mediterranean coast and Andalusia Atlantic coast). ** indicates significance at P < 0.001.

The NW Mediterranean experienced a marked and synchronous decline in *C. gallina* landings across all four AACCs during the late 1990s and early 2000s (Fig. 2A). Landings peaked at 1,580 tonnes in 1995 and then collapsed by more than 80%, falling to 46.8 tonnes by 2004. A slight recovery occurred in 2005 (126.3 tonnes), after which landings declined gradually, reaching zero by 2021. In recent years (2023–2024), catches have resumed only in Catalonia and remain below 8 tonnes per year. Landings of *D. trunculus* reached their maximum in 1993, at 1,137 tonnes (Fig. 2B), before dropping sharply to 115 tonnes by 1999. A modest recovery occurred in the early 2000s, with a secondary peak in 2004 (612 tonnes), followed by a continuous decline throughout the 2010s and very low levels by the early 2020s. For *C. chione*, landings peaked in 1994 at 302 tonnes (Fig. 2C), but the fishery collapsed by 1998. Since the early 2000s, reported catches have remained close to zero, with no signs of recovery through 2024. Strong positive and significant correlations were detected among the three species in the NW Mediterranean: *C. gallina* and *D. trunculus* (r = 0.866, P < 0.001), *C. gallina* and *C. chione* (r = 0.847, P < 0.001), and *D. trunculus* and *C. chione* (r = 0.849, P < 0.001).

**Figure 2.**
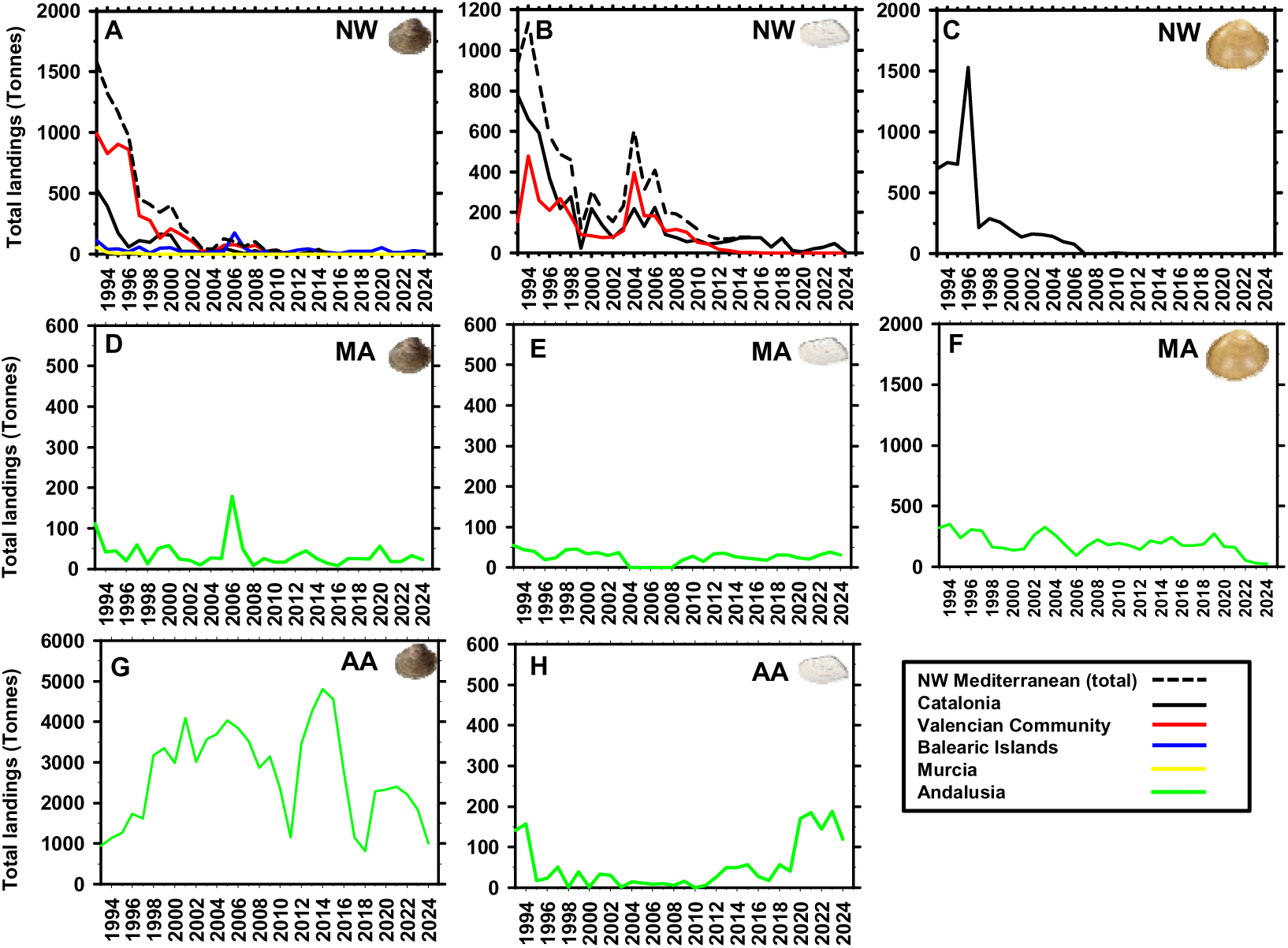
Annual landings (tonnes) of the three main commercial clam species: *(A, D, G) Chamelea gallina*; *(B, E, H) Donax trunculus*; and *(C, F) Callista chione*, across three geographic subregions: (A-C) Northwest Mediterranean (Catalonia, Valencian Community, Balearic Islands, and Murcia); (D-F) Mediterranean Andalusia (Alboran Sea); and (G-H) Atlantic Andalusia (Gulf of Cadiz). Coloured lines indicate landings by AACC: black for Catalonia, red for Valencian Community, blue for Balearic Islands, yellow for Murcia, and green for Andalusia. The dashed black line in panels A and B represents the aggregated landings from all Northwestern Mediterranean AACC combined.

On the Mediterranean coast of Andalusia, *C. gallina* landings showed a fluctuating pattern between 1993 and 2024 (Fig. 2D). Catches declined from 111.2 tonnes in 1993 to 8.8 tonnes in 2003, then rebounded sharply to 178 tonnes in 2006. This recovery was short-lived, however, and landings fell again to 7.4 tonnes in 2008. A moderate rebound followed between 2009 and 2015, with annual catches fluctuating between 14 and 44 tonnes. After a further decline in 2016 (7.5 tonnes), landings gradually increased from 2017 to 2024, stabilising between 18 and 57 tonnes. This irregular trajectory may reflect unstable stock dynamics, shifts in fishing pressure, or regulatory changes. *D. trunculus* also showed a variable trend over the study period (Fig. 2E). Landings peaked in 1993 at 55 tonnes, then declined progressively to a minimum of 0.56 tonnes in 2005. A brief recovery occurred in 2006, but landings subsequently fell to zero in 2007, likely reflecting the temporary cessation of fishing activity rather than the absence of the resource, as reported by local fishers and management records. From 2008 onwards, catches stabilised at relatively low levels, fluctuating between 18 and 35 tonnes per year with limited interannual variability. In contrast, *C. chione* landings remained relatively stable along the Mediterranean coast of Andalusia from 1993 to 2021 (Fig. 2F), ranging from 130 to 351 tonnes per year, apart from a slight dip in 2006 (93 tonnes). This trend shifted recently: since 2022, reported catches have declined markedly and have not exceeded 60 tonnes per year through 2024. No significant correlations were detected among the three species on the Mediterranean coast of Andalusia (i.e., *C. gallina* and *D. trunculus*, r = 0.162, P = 0.374; *C. gallina* and *C. chione*, r = −0.084, P = 0.645; and *D. trunculus* and *C. chione*, r = 0.187, P = 0.645).

On the Atlantic coast of Andalusia, *C. gallina* landings displayed the highest productivity of the three management areas (Fig. 2G) and followed a pronounced boom-and-bust pattern. After increasing from 1993 to 1998, catches remained relatively stable from 1998 to 2009, ranging between 2,800 and 4,100 tonnes per year. A sharp decline then occurred in 2010, when landings dropped to 1,153 tonnes. This was followed by a strong recovery from 2012 to 2016, with annual catches fluctuating between 2,700 and 4,800 tonnes. In subsequent years, the fishery showed moderate variation: a decline in 2017–2018 (1,146 and 810 tonnes, respectively), a partial recovery in 2019–2022 (2,200–2,400 tonnes), and another decline in 2023 and 2024 (1,840 and 1,025 tonnes, respectively). This trajectory indicates a highly dynamic fishery, probably shaped by both environmental and regulatory variability. In the same region, *D. trunculus* landings showed marked interannual variability throughout the study period (Fig. 2H), with three broad phases. The first phase (1993–2003) showed a pronounced decline, from 157.5 tonnes in 1993 to just 0.6 tonnes in 2003. The second phase (2003–2011) was characterised by persistently low landings, generally below 15 tonnes per year. In contrast, the third phase (2011–2024) showed a progressive recovery despite interannual fluctuations, with catches ranging from 60 to 110 tonnes annually and peaking at 186 tonnes in 2023. No significant correlation was detected between *D. trunculus* and *C. gallina* landings in this area (r = 0.307, P = 0.086, N = 32). Unlike the Mediterranean coast, *C. chione* is not commercially harvested in this area.

### 3.2. Environmental variability and correlations with landings

Environmental variability showed distinct temporal patterns across the three study regions (Fig. S1). SST anomalies generally increased during the later years of the time series, whereas SSS and chlorophyll-a anomalies showed greater interannual variability. Pearson correlation analyses between annual landings and environmental variables revealed generally weak and inconsistent relationships across species and regions (Table S1). Significant correlations were detected only in a few cases. Negative correlations with SST were observed for *C. chione* in the Alboran Sea and for *C. gallina* and *D. trunculus* in the NW Mediterranean. A positive correlation between chlorophyll-a anomalies and *C. chione* landings was detected in the NW Mediterranean. Overall, no consistent environmental pattern was identified that could explain the contrasting trajectories of clam fisheries across regions.

### 3.3. Governance and management

The governance of small-scale fisheries (SSF), including clam fisheries, has been progressively decentralised in Spain since the early 1980s, following the approval of national decrees delegating responsibilities to the regional governments of the AACCs (e.g., Catalonia: RD 1646/1980; the Balearic Islands: RD 3540/1981; the Valencian Community: RD 3533/1981; Murcia: RD 4190/1982; and Andalusia: RD 3506/1983). As a result, each AACC has independently regulated clam fisheries within its territorial waters, producing a patchwork of management approaches with no interregional coordination; the marked differences in minimum legal size (MLS) for *C. gallina* and *D. trunculus* among AACCs provide one example (Fig. 3). During the 1980s and 1990s, AACCs legislation established basic appropriation rules, including restrictions on user access (e.g., shellfish-harvesting licences), gear specifications (e.g., mesh sizes and number of dredges), MLS, mandatory screening of specimens at fishers’ guilds, and total allowable catch (TAC) limits. These measures were progressively reinforced in the 2000s in response to declining catches. The 2010s marked a turning point with the adoption of formal fishery-specific management plans, particularly in response to EU requirements (Council Regulation (EC) 1967/2006 and Regulation (EU) No 1380/2013). Since the 2020s, these plans have continued to be updated. Nonetheless, despite divergent governance trajectories across regions and substantial regulatory effort, most fishing grounds have experienced pronounced declines or prolonged closures, highlighting the persistent challenge of achieving sustainable and adaptive governance in clam fisheries (Table 2; Fig. 4).

**Figure 3.**
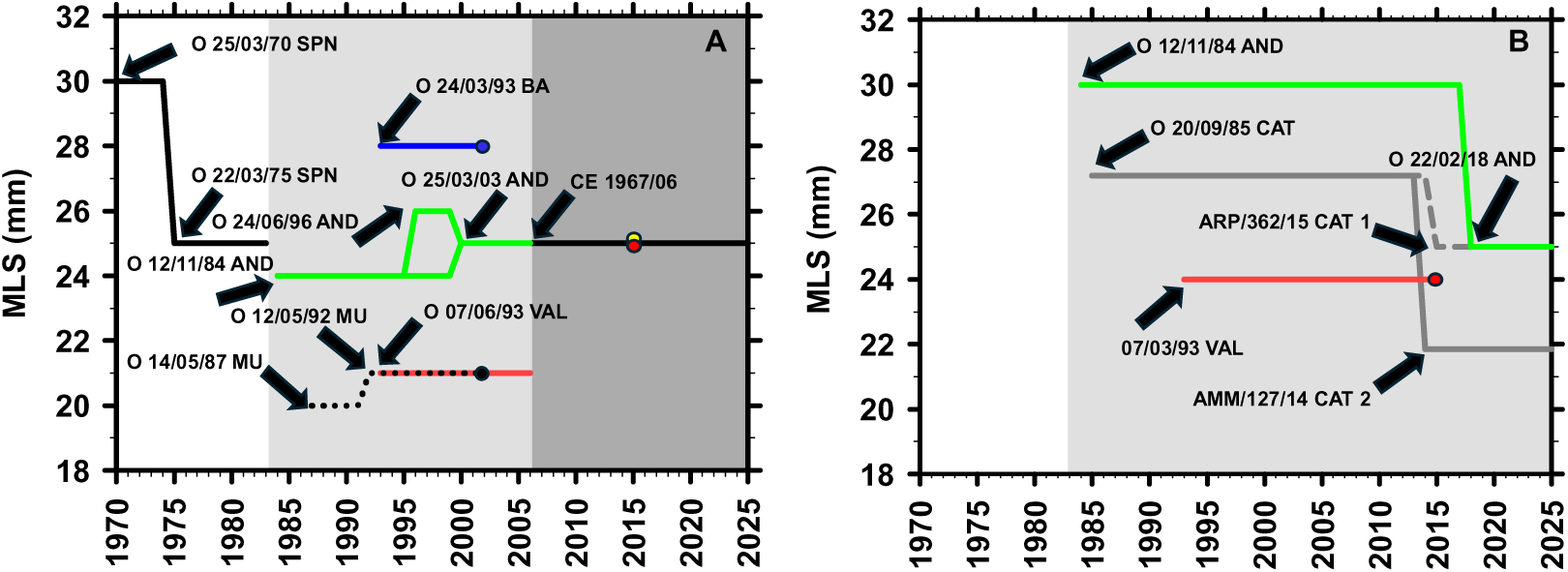
Evolution of the MLS in Spain. **(A)** *C. gallina*: The solid black line indicates the national MLS set by the central government between 1970 and 1983. After the decentralisation of fisheries competences (1980-1983), AACCs established their own MLS regulations. Red line: Valencian Community; dotted black line: Murcia; blue line: Balearic Islands; green line: Andalusia. From 2006 onwards, MLS was harmonised across Spain by the European regulation EC 1967/06. Catalonia is absent from the series because no MLS was defined until 2006. In the Valencian Community, Order 07/06/1993 did not specify an MLS but required sieving with a 19-mm circular mesh, which according to [38] corresponds to specimens larger than 21 mm. In the Balearic Islands, Order 24/02/1993 established a shell thickness >15 mm, equivalent to a total length of 28 mm (our data). In Andalusia, between 1996 and 2003, two different minimum sizes coexisted: 26 mm for hydraulic dredges and 24 mm for the remaining fishing gears. Dots indicate the year when fisheries were closed: blue for Balearic Islands, black for Murcia, red for Valencia, yellow for Catalonia. Letters inside the figure identify the regulation establishing each MLS (SPN, Spain; VAL, Valencian Community; BA, Balearic Islands; MU, Murcia; AND, Andalusia; CE, European Community). **(B)** *D. trunculus*: Grey solid line shows Catalonia with hand-operated dredges; grey dashed line shows Catalonia with mechanised dredges; red line: Valencian Community; green line: Andalusia. The red dot marks the closure of the fishery in Valencia. In this case, Order 07/06/1993 required sieving with a 14 mm circular mesh, corresponding to specimens larger than 24 mm [38]. In Catalonia, Order AAM/127/2014 set a shell thickness of 7 mm, equivalent to a total length of 21.9 mm (our data).

**Table 2.**
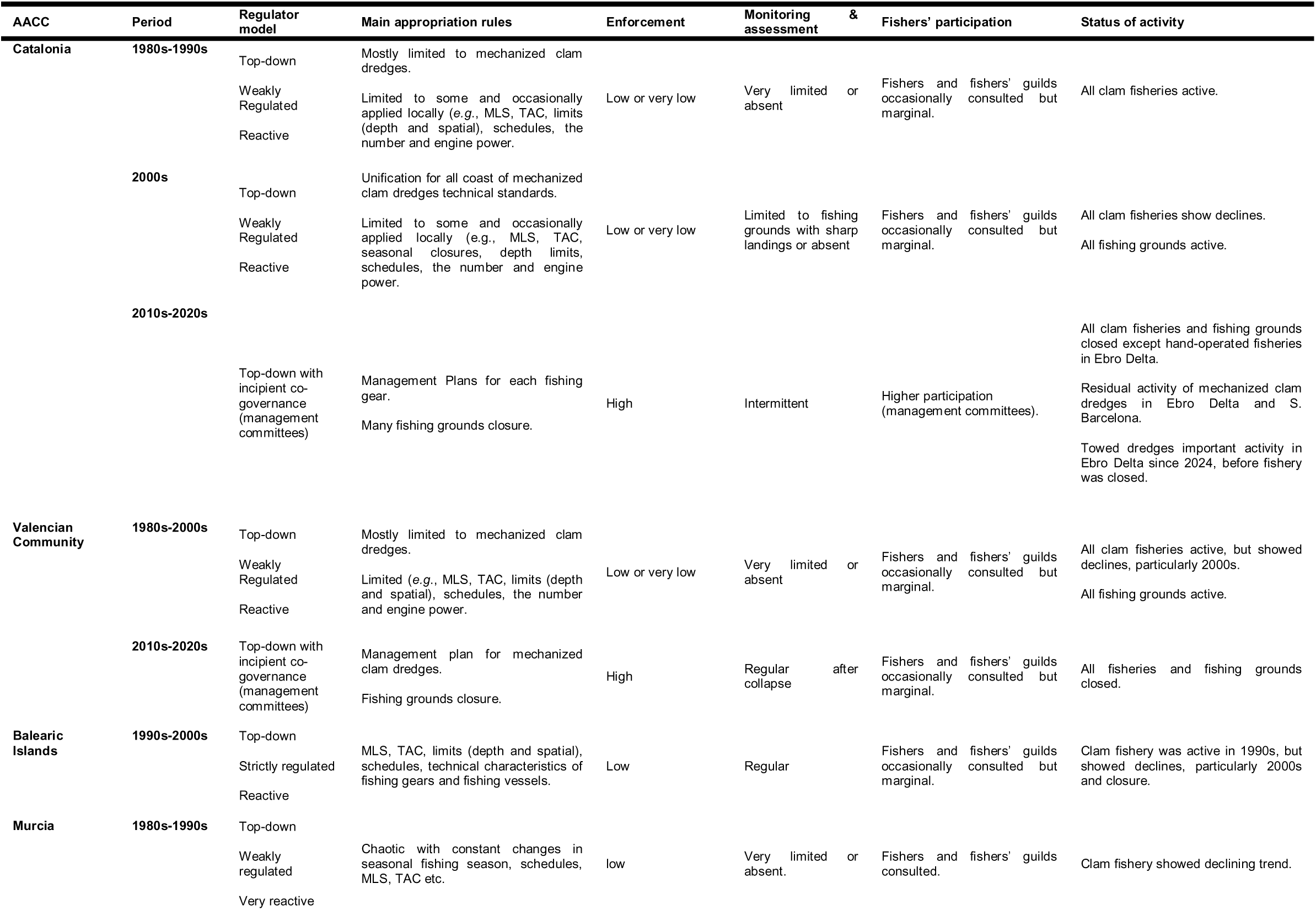
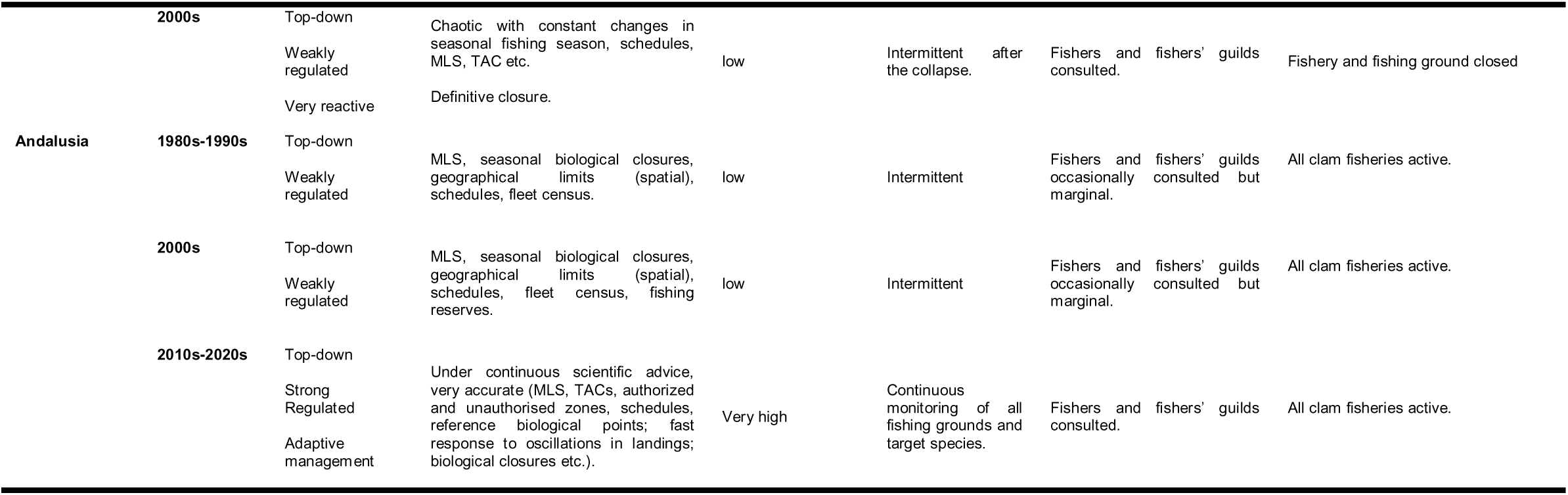
Governance and management of clam fisheries in Spanish AACC (1980s-2020s). Summary of regulatory models, appropriation rules, enforcement, monitoring and assessment, fishers’ participation and activity status across regions.

**Figure 4.**
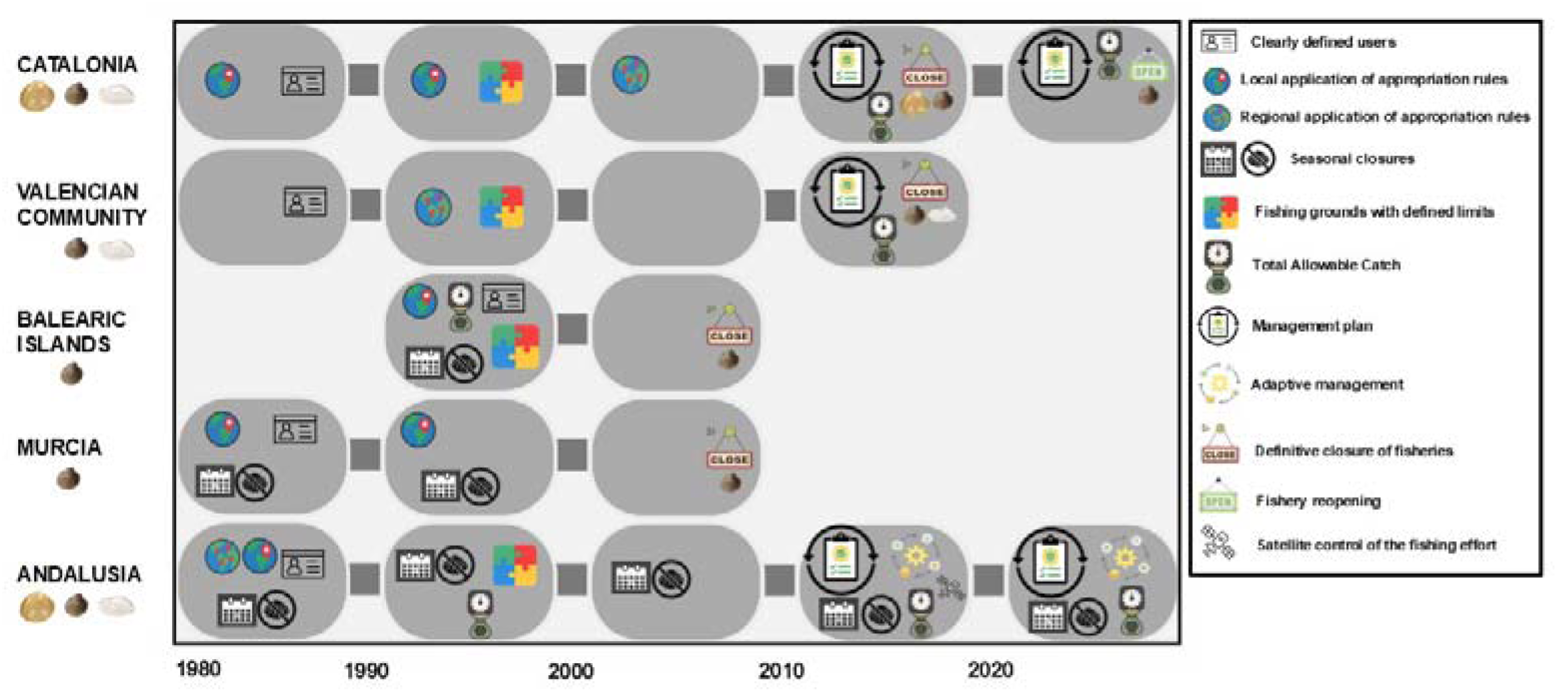
Chronological synthesis of the main regulatory and management measures affecting clam fisheries in Spain by autonomous communities (AACC), grouped by decade. The figure illustrates the transition from basic appropriation rules during the 1980s-2000s to more structured management frameworks, monitoring systems, and adaptive management approaches during the 2010s-2020s. Symbols represent the following categories: restricted access (licensing of shellfish harvesters), local fishing-ground regulations, regional regulations, seasonal closures, spatially defined fishing grounds, Total Allowable Catch (TAC), management plans, adaptive management measures, definitive fishery closures, fishery reopening, and satellite-based monitoring of fishing effort.

#### Clam fisheries in the Northwestern Mediterranean Sea

Clam fisheries in Catalonia historically relied on three main gears: winch-operated mechanised dredges (*C. gallina*, *D. trunculus*, and *C. chione*), vessel-towed dredges (mainly *C. gallina*), and hand-operated dredges used by shore-based shellfishers (*D. trunculus*). This diversity created multiple user groups and access regimes shaped by both ecological conditions and evolving institutional rules.

The first regulatory framework emerged in the mid-1980s, when Order 20/09/1985 established an MLS for *D. trunculus*, followed by Decree 9/1987, which created separate licensing schemes for mechanised dredges and hand-operated gears. During the late 1980s and 1990s, additional orders defined effort restrictions, sub-zoning of fishing grounds, and daily quotas, although enforcement was uneven and several areas remained poorly regulated. Conflicts between mechanised and hand dredgers, especially in the Ebro Delta, were frequent, while vessel-towed dredges targeting *C. gallina* operated with limited legal coverage.

In the 2000s, Catalonia attempted institutional consolidation. Order AAR/143/2007 unified technical standards for mechanised dredges, but compliance with gear specifications was poor and landings continued to decline. Reactive measures, including short-term closures (*e.g*., AAR/60/2008), were inconsistently applied and often reversed after disputes with other sectors.

The adoption of EU-driven management plans in the 2010s marked a significant shift. Orders ARP/362/2015 and ARP/122/2020 established gear-specific frameworks for mechanised dredges; Orders ARP/114/2018 and ARP/188/2020 did the same for vessel-towed dredges; and Orders AAM/127/2014, ARP/256/2016, and ARP/182/2019 regulated hand-operated dredges. These plans introduced defined fishing grounds, technical standards, and participatory committees involving fishers, scientists, and administrators, although decision-making remained in administrative hands. By the early 2020s, nearly all mechanised dredging grounds had been closed, except for residual activity in the Ebro Delta and southern Barcelona. A partial reopening occurred in 2024, when *C. gallina* was again authorised for mechanised and towed dredges in specific grounds (see the Supplementary Material for the full regulatory chronology). Despite these advances, governance in Catalonia remains fragmented and inconsistently enforced, with limited integration of ecological knowledge and weak adaptive capacity. Current activity is residual, and clam fisheries continue to operate under precarious conditions.

Clam fisheries in the Valencian Community have historically relied on two distinct gears: mechanised dredges operated from vessels, targeting *C. gallina* and *D. trunculus*, and hand-operated dredges used by shore-based shellfishers for *D. trunculus*. These two modalities have long differed in their operational dynamics, access regimes, and levels of institutional oversight.

The first legal framework emerged during the late 1980s and 1990s. Order 15/09/1988 introduced a clam fishing licence, and Decree 67/1996 consolidated this system by formally distinguishing mechanised dredges from hand-operated dredges, thereby creating separate user groups. At the same time, the Resolution of 02/06/1993 classified production areas, and the Order of 07/06/1993 introduced technical limits on gear (maximum mouth widths of 75 cm for mechanised and 85 cm for hand-operated dredges), mesh sizes (19 mm for *C. gallina*, 14 mm for *D. trunculus*), and daily catch quotas. For hand-gatherers, permits were capped at 25 per guild. These rules clarified appropriation regimes but were poorly enforced and disconnected from ecological indicators, leaving the fishery highly vulnerable to overexploitation. Throughout the 2000s, no substantial new measures were introduced despite steep declines in landings, mirroring the collapse seen in Catalonia.

Stronger regulations arrived only in the 2010s. Decree 94/2013 updated the legal framework, followed by Decree 62/2016, which established a comprehensive management plan covering eligibility rules, fishing schedules, technical specifications, MLS, and total allowable catches (TACs). In parallel, closures were implemented: Resolution 09/08/2013 imposed indefinite bans on three fishing grounds (Patacona-Cabanyal, Molinell-Deveses, and Tavernes). Resolution 03/06/2015 extended the closure to the entire Valencian coast for mechanised dredges and to most areas for hand-operated dredges, effectively shutting down the fishery. Resolution 18/05/2017 further extended the ban on hand-operated dredges targeting *D. trunculus* to the remaining areas of the Valencian coast. The initial closed areas were intended to act as larval reserve zones, although no empirical evidence supported their ecological effectiveness. In practice, by 2015 the closure was almost complete because hand-gatherers operating outside the prohibited areas lacked authorised depuration facilities, resulting in irregular activity. Follow-up commissions involving guilds, scientists, and administrators were also established, representing a first step towards co-governance, although their role remained consultative and ultimate authority continued to reside with the administration (see the Supplementary Material for the full regulatory chronology). Today, clam fisheries in the Valencian Community remain inactive, with indefinite closures in force and little evidence of recovery despite continued monitoring.

The clam fishery in the Balearic Islands was formally regulated in 1993, following the discovery of a productive ground for striped venus clams (*C. gallina*) off Palma Bay. From the outset, only vessel-based mechanised dredges were permitted. Order 24/02/1993 established a mandatory licensing system, set daily catch limits based on crew size (15-20 kg for one fisher and 40-60 kg for three), and imposed a total allowable catch of 70 tonnes for the entire fleet. Each vessel was limited to two dredges, and working hours were initially capped at nine, later reduced to seven. Technical specifications for dredges and onboard mesh sizes were also defined. Furthermore, all catches were required to be inspected by local guilds before sale to ensure compliance with minimum size requirements.

Throughout the 1990s, regulations were frequently updated to refine appropriation rules. Annual adjustments included Order 31/10/1994 and a series of resolutions issued in 1995, 1996, 1997, and 1998, which modified schedules, quotas, and technical limits. Scientific monitoring complemented this adaptive approach, but concerns about declining landings and early signs of resource depletion persisted. In 2002, the regional government closed the Palma Bay fishery as a precaution. The closure was later extended without a recovery plan, regular monitoring, or reopening rules. A 2003 amendment to production zones was introduced but failed to establish specific provisions for clams. Consequently, the fishery has remained closed, lacking an effective framework for revival (see the Supplementary Material for the full regulatory chronology). Despite relatively stringent appropriation rules and repeated regulatory updates, governance in the Balearic Islands remained highly centralised and rigid, with no participatory structures and little adaptive capacity. This case illustrates how a management approach centred on control and restriction, without meaningful stakeholder involvement, was insufficient to prevent collapse or support resilience in small-scale clam fisheries.

The case of Murcia is particularly noteworthy within the Spanish context because regional authorities inherited, in the 1980s, a clam fishery that was already experiencing a marked decline in landings. From the outset, management was reactive and often improvised, frequently alternating between closures, adjustments to minimum size limits, and restrictions on fishing effort in an attempt to balance resource protection with fishers’ demands.

The Resolution of 09/04/1984 introduced a minimum landing size (MLS) of 18 mm together with a seasonal closure; however, this measure was swiftly repealed in 1985. A new regulatory cycle began with the Order of 09/04/1986, which reinstated the closure, increased the MLS to 20 mm, and imposed rules on users, dredge specifications, and fishing schedules. In subsequent years, frequent changes highlighted the absence of strategic planning. The Order of 09/09/1987 extended the fishing season, whereas the Order of 12/05/1988 restricted it to June, reduced fishing hours, and introduced daily total allowable catches (TACs). Thereafter, regulations alternated between extending and limiting fishing periods (e.g., Orders of 10/04/1990, 28/05/1991, and 12/06/1992), while progressively raising the MLS to 21 mm and restricting the number of dredges per vessel. At the same time, the Resolution of 30/06/1995 closed one of the main fishing grounds (Cabos Negrete-Palos), despite having been repealed by a conflicting measure the previous year. Despite this constant regulatory fluctuation, landings continued to decline throughout the 1990s. Ultimately, the Order of 16/06/2002 permanently closed all clam fishing grounds in Murcia, effectively shutting down the fishery. Scientific surveys conducted thereafter revealed no signs of recovery, and the closure remains in effect today (see the Supplementary Material for the complete regulatory chronology). In summary, Murcia exemplifies a highly reactive governance system characterised by frequent short-term regulatory adjustments—covering MLS, fishing seasons, gear limits, and catch quotas—without a coherent long-term strategy. This unstable framework failed to halt overexploitation or promote adaptive management and ultimately culminated in fishery collapse.

#### Clam Fisheries in Mediterranean and Atlantic Coast of Andalusia

Clam fisheries in Andalusia have developed under distinctly different conditions along the Mediterranean and Atlantic coasts. In the Mediterranean, the fishery primarily depends on vessel-based mechanised dredges operated by winches and targeting species such as *D. trunculus*, *C. gallina*, and *C. chione*. In the Atlantic, by contrast, *D. trunculus* is harvested with hand-operated dredges, whereas *C. gallina* is targeted with engine-powered towing and mechanised hydraulic dredges.

The initial rules for appropriation were established in the mid-1980s. Order 16/02/1984 set minimum legal sizes (MLS) of 24 mm for *C. gallina*, 30 mm for *D. trunculus*, and 60 mm for *C. chione*, as well as implementing seasonal closures. A shellfishing licence was introduced in Order 19/11/1984, while the fleet underwent its first census as mandated by Order 23/11/1986. Throughout the 1990s, geographic boundaries were defined (Orders 25/04/1993, 22/12/1998, 09/11/1999), and updates to the census were conducted (Order 24/10/1994). In the Atlantic, dredge fisheries gradually came under regulation: Order 24/06/1996 established technical standards for towed dredges, and Order 25/03/1999 governed hydraulic dredges, including stipulations on depth limits, catch quotas, and MLS (ranging from 24 to 26 mm). During the 2000s, the regulatory framework saw only minor updates, notably with Order 25/03/2003, which raised the MLS for *C. gallina* to 25 mm. A significant development was the creation of the Guadalquivir estuary reserve (Orders 16/06/2004 and 11/01/2005), which introduced zoning and distinct regulations for hydraulic dredges, towed dredges, and hand-gatherers.

The 2010s marked a major shift in shellfishing regulation. Decree 387/2010 and Decree 99/2015 distinguished between on-foot and vessel-based shellfishing, establishing specific licensing schemes for each. In the Atlantic, following a collapse in landings, the Guadalquivir *C. gallina* grounds were subject to a precautionary closure (Order 16/12/2010), followed by temporary closures (Orders 30/11/2016 and 29/06/2017). The latter also introduced a comprehensive management framework incorporating quotas, satellite tracking, and monitoring. At the same time, working hours and minimum landing sizes (MLS) were standardised through Orders 26/02/2013, 13/06/2013, 23/11/2017, and 22/02/2018. In the Mediterranean, gear-specific management plans were introduced (Order 24/03/2014; Order 27/12/2019), including quotas, vessel monitoring, and sanctioning systems (see the Supplementary Material for the full regulatory chronology).

From 2020 onwards, Andalusia has developed one of the most advanced management systems in Spain. A dedicated order dated 06/04/2020 established a management plan for *C. gallina* in the Atlantic, followed by annual fishing plans (Resolutions 22/07/2021, 04/11/2022, and 13/07/2023) and precautionary closures when thresholds were exceeded (Resolutions 08/02/2021, 09/11/2022, and 16/03/2023). For *D. trunculus*, Order 25/05/2020 regulated hand-operated dredges in the Atlantic, later consolidated in a management plan under Order 20/07/2023 and likewise enforced through temporary closures (Resolution 05/10/2023). In the Mediterranean, adaptive plans were implemented, with closures triggered by scientific assessments (Resolutions 05/12/2024 and 17/02/2025 for *D. trunculus*, and 18/10/2024, 27/02/2025, and 26/06/2025 for *C. gallina*) (see the Supplementary Material for the full regulatory chronology). Overall, Andalusia has progressively established a comprehensive, adaptive governance system, where scientific monitoring, satellite tracking, annual plans, and rapid closures are directly linked to stock assessments and management decisions. In contrast to other regions, Andalusia now serves as a benchmark for adaptive small-scale fishery (SSF) management in Spain.

## 4. Discussion

This study shows that small-scale clam fisheries have followed markedly different long-term trajectories across Spanish regions. These contrasting trends reflect a combination of environmental conditions, fishing pressure, and governance and management frameworks. The three regions analysed here—the NW Mediterranean, the Alboran Sea, and the Gulf of Cádiz—differ substantially in their hydrographic and trophic conditions. The NW Mediterranean is generally more oligotrophic, whereas the Alboran Sea and the Atlantic coast of Andalusia are comparatively more productive because of Atlantic inflow and higher nutrient availability [45–47]. Despite historically intense fishing pressure across all three regions, their fisheries have evolved differently. Whereas the NW Mediterranean has experienced repeated collapses or prolonged declines, fisheries in the Alboran Sea and the Gulf of Cádiz have shown greater resilience. These contrasting trajectories are consistent with the joint influence of ecological dynamics and governance structures on fishery outcomes. They also coincide with substantial differences in the timing and intensity of management interventions across regions, suggesting that long-term fishery dynamics are shaped by the combined influence of environmental variability and governance responses. Commercial clam populations commonly exhibit strong interannual variability in landings [6,7,48,49]. Their population dynamics are influenced not only by fishing pressure but also by environmental conditions affecting spawning, recruitment success, juvenile survival, condition index, abundance, and biomass [7,18,24,50,51]. However, the environmental variables analysed in this study did not show a consistent relationship with the long-term decline in clam landings across regions, suggesting that environmental forcing alone cannot explain the contrasting trajectories documented here.

Clams may also experience occasional mass mortality events that abruptly reduce stock biomass and strongly affect subsequent landings [51,52]. Such ecological variability is therefore an inherent feature of many clam fisheries and requires management systems based on continuous monitoring and adaptive regulation capable of responding to rapid changes in stock dynamics. To our knowledge, no long-term ecological studies have documented recurrent recruitment failures, large-scale mortality events, or sustained biomass declines for the clam fisheries analysed here. Most available information derives from landing statistics rather than direct stock assessments. In addition, high historical landings may sometimes reflect periods of intensive exploitation of previously abundant stocks rather than high underlying ecosystem productivity, as may have occurred in parts of the NW Mediterranean during the 1990s.

Declines in landings or stock biomass of several commercially exploited clam species have also been reported in nearby regions outside the study area during recent decades. For example, declines in *D. trunculus* have been reported in the Camargue (France, NW Mediterranean) [53,54], in Galicia (NW Spain) [55], and in Agadir (Atlantic Morocco) [56]. Similar declines have been documented for *C. gallina* and *Ensis minor* in the Adriatic Sea [57,58], as well as for *C. chione* along the Mediterranean coast of Morocco [59]. These observations suggest that the declines identified here are not unique to the Spanish fisheries analysed in this study but may reflect broader regional patterns affecting Mediterranean and adjacent Atlantic clam fisheries. Such patterns indicate that commercially exploited clam species, often subject to intense fishing pressure and environmental variability, may respond similarly across different regions. However, the contrasting trajectories observed among Spanish regions also suggest that environmental variability alone cannot explain the differences in fishery outcomes. Understanding the mechanisms that facilitate connectivity among fishing grounds is therefore essential for developing sustainable management strategies for commercially exploited species [60]. The dispersal of pelagic larvae can connect spatially separated subpopulations and potentially generate metapopulation structures [61] The duration of the pelagic larval stage determines how long larvae remain in the plankton and therefore their potential dispersal distance, which may extend over considerable spatial scales through the action of currents and other physical processes [62,63]. For the species analysed here, the pelagic larval stage lasts approximately 15–30 days for *C. gallina*, around 40–60 days for *D. trunculus*, and 32–39 days for *C. chione* [64–67]. Along the Iberian coast, several oceanographic discontinuities influence larval transport and population connectivity. Among the most prominent are the Strait of Gibraltar and the Almería-Oran Front (AOF), both recognised as persistent biogeographic and genetic breaks across multiple marine taxa [68]. The AOF separates the oligotrophic Balearic Sea in the Northwestern Mediterranean from the more productive Alboran Sea [69], whereas the Strait of Gibraltar is the main exchange zone between Atlantic and Mediterranean waters [70]. The strong correlations observed among clam landings within the Northwestern Mediterranean suggest a relatively high degree of connectivity within this subregion. The absence of major hydrodynamic barriers may facilitate larval exchange among fishing grounds located in different Spanish AACCs. By contrast, the lack of correlation between the Northwestern Mediterranean, the Alboran Sea, and the Atlantic coast of Andalusia is consistent with oceanographic and genetic evidence indicating that the Almería-Oran Front and the Strait of Gibraltar may act as barriers to connectivity for several marine taxa, including clams [35]. These results suggest that ecological connectivity may extend beyond administrative boundaries, highlighting the importance of coordination among regional fisheries authorities to ensure coherent management of shared clam stocks.

These ecological patterns highlight a potential mismatch between biological population structure and the administrative scales at which fisheries are managed. Differences in regulatory design and enforcement capacity appear to have amplified the contrasting ecological trajectories observed across subregions. Where management has been largely reactive—intervening only after clear declines in landings—and guided by a short-term perspective, recovery of clam stocks has often been limited or absent, even in areas with moderate environmental productivity, such as the Ebro Delta in Catalonia. This type of crisis-driven intervention resembles the “event-based change” described in organisational theory: short-term, reactive responses to unpredictable shocks that disrupt established routines [71]. Similar dynamics have been described by Pedroza and Salas [72] for the Yucatán fisheries, where governance was dominated by event-based, reactive strategies and struggled to achieve sustainability. While such responses may alleviate acute problems, they rarely address the structural vulnerabilities that predispose fisheries to collapse. By contrast, proactive approaches are more closely aligned with the notion of “time-paced evolution”, whereby adaptive adjustments are introduced continuously and deliberately over time to build resilience rather than merely react to crises [73]. This concept strongly resonates with adaptive fisheries management, which seeks to move from reactive responses to structured systems of continuous learning [74–76]. Adaptive management emphasises learning from both successes and failures and incorporating new information into decision-making, thereby reducing uncertainty and supporting resilience. It is particularly recommended for fisheries with limited data and institutional capacity [77,78]. In our case study, Andalusia provides the strongest example of this approach over the last two decades. The region has implemented annual effort limits, dynamic closures, and gear-specific regulations, complemented by continuous resource assessments (e.g., spatially detailed monitoring) and rigorous compliance controls. The gradual incorporation of digital monitoring tools and decision-support systems has further strengthened this adaptive framework, including electronic catch reporting, real-time quota tracking, and adaptive spatial management. Together, these tools have enhanced both transparency and responsiveness.

Beyond these institutional and technological advances, the spatial context of the fishery also plays an important role. The location of the main clam fishery within Doñana National Park [79], the Marine Protected Area of the Guadalquivir estuary [80], and the Fishing Reserve established at the mouth of the Guadalquivir River by the Order of 16 June 2004 constitutes an important element of control and surveillance aimed at preventing excessive fishing pressure. Its position within this highly protected area implies stricter monitoring protocols, stronger enforcement capacity, and greater institutional attention, which together reduce the likelihood of illegal, unreported, and unregulated (IUU) fishing. This setting also creates a governance framework in which conservation objectives and fisheries management are more closely aligned, potentially leading to higher compliance and more effective limitation of fishing effort. It may therefore represent a distinctive feature of these fisheries relative to the other stocks examined here, as the regulatory and ecological constraints associated with the park’s buffer zone provide an additional layer of protection beyond conventional governance frameworks.

Fisheries are often managed according to administrative or political jurisdictions rather than ecological scales [81,82]. This practice frequently generates spatial mismatches, particularly when jurisdictional boundaries are too narrow to capture the dynamics of exploited populations. Such mismatches are a major threat to fishery sustainability [83]. Because such misalignments can reduce productivity and promote the overexploitation of distinct subpopulations [84–86], they have been documented in many commercially exploited fish species, both pelagic and demersal, [83,87], but they are equally evident in sedentary or low-mobility invertebrates [88,89]. The stalked barnacle (*Pollicipes pollicipes*) fisheries in Spain, Portugal, and France provide one example: administrative management plans divide populations that are biologically connected [88]. The sea urchin (*Paracentrotus lividus*) fishery in Galicia provides another, where local management units do not account for genetic connectivity among subpopulations, increasing the risk of depletion of local populations [90]. The same problem is evident in Spanish clam fisheries, where regional management by AACCs does not align with the metapopulation structure of the target species in the Northwestern Mediterranean. This misalignment reinforces stock vulnerability and limits the effectiveness of recovery measures.

Despite their considerable social and ecological importance, SSFs have historically received limited attention in the EU. For decades, they have been subject to weaker management, monitoring, and regulatory frameworks than their large-scale counterparts, largely because of their comparatively lower catches and economic value [13,91]. However, SSFs play a key role in employment and can represent an important source of local economic activity. Beyond their economic contribution, they strengthen attachment to place and contribute to social cohesion and stability in rural and peripheral communities [13]. Spanish clam fisheries illustrate this pattern clearly, especially in the NW Mediterranean, where they were often poorly regulated or virtually unmanaged for long periods by the AACCs. In Catalonia, the collapse of *C. chione* in the Maresme region after 2007 led to the *de facto* abandonment of the fishery, without monitoring or recovery plans. In the Ebro Delta, clam dredging operated under minimal regulation until the 2010s. In the Valencian Community, much of the 1990s and 2000s resembled a quasi-open-access regime, with exploitation of *C. gallina* and *D. trunculus* occurring under nominal restrictions. This reflected both limited administrative capacity and the perception that these small-scale activities did not justify sustained investment in management. Regulations were generally limited to a small set of appropriation rules—such as user restrictions, MLS, seasonal closures, and gear restrictions—while systematic monitoring and scientific stock assessment were largely absent or inadequate. This scenario is common in SSF, where costs and logistical challenges are comparatively high[13,92]. Since the transfer of fisheries-management responsibilities from the central government to the AACCs in the 1980s, governance of Spanish clam fisheries has consistently followed a top-down model across all regions analysed here.

In this traditional regulatory model, power remains concentrated in government agencies, and decisions are often made without sufficient consideration of the realities faced by fishing communities. This mirrors a broader European pattern in which SSFs have been marginalised under policy frameworks designed primarily for industrial fleets, with little adaptation to the socio-ecological specificities of artisanal sectors [93]. Under this model, the involvement of fishers and Cofradías (fishers’ guilds) in governance was generally limited to consultation and gained visibility only once stocks had entered a critical state of decline. In Spain, fishers’ guilds are the main institutions that organise and represent small-scale fishers and shellfish harvesters, although their membership often extends to operators in more industrialised fleets. These guilds function as hybrid entities with both public and private dimensions [94], serving as advisory and collaborative counterparts to regional administrations in fisheries governance [95] in Spain. The limited inclusion of fishers and guilds in governance has undoubtedly contributed to recurrent stock depletion and, ultimately, to the collapse and closure of many clam fisheries, particularly in less productive areas such as the NW Mediterranean. Conventional top-down management approaches are increasingly viewed as inadequate, highlighting the need for governance models that integrate both ecological sustainability and social dimensions [96].The European Commission [97] likewise stressed the need to strengthen fishers’ involvement in decision-making to improve institutional legitimacy and increase compliance with regulatory measures.

In recent years, European research has paid increasing attention to decentralisation, the transfer of responsibilities, and the active involvement of fishers in management frameworks [98–100]. Participatory co-governance models, also known as fisheries co-management, encompass a broad spectrum of institutional arrangements in which management responsibilities are shared between state administrations and resource users. These systems range from instrumental forms, in which participation mainly facilitates the implementation of predefined rules, to empowering forms, in which fishers genuinely contribute to the development of regulations [101,102]. Evidence from many regions shows that co-governance can improve compliance, incorporate fishers’ ecological knowledge into decision-making, and strengthen the legitimacy and resilience of management systems [103]. The Asturian stalked barnacle fishery is one European example, with Territorial Use Rights for Fisheries (TURFs) managed by Cofradías demonstrating long-term adaptive capacity[104]. Similar experiences in Chile and northwestern Mexico show that co-management works best when enabling legislation exists, rights are clearly defined, and institutions are stable [101,105]. These examples suggest that Spanish clam fisheries could benefit from moving beyond symbolic consultation towards genuine co-governance, in which Cofradías and fishing communities play a formal role in planning, oversight, and enforcement.

Of the AACCs analysed, Catalonia is the only region to have implemented a governance framework inspired by co-management, as established in Decree 118/2018 on the professional fisheries governance model. However, clam fisheries are not explicitly included within this framework. Instead, gear-specific monitoring committees—covering mechanised dredges, hand-operated dredges, and tow dredges—were introduced as putative co-management bodies. In practice, these committees remain primarily consultative: although fishers, scientists, and managers meet regularly, their conclusions are advisory only, and final authority remains with the administration. This represents a limited form of participation rather than genuine co-management and does not meet the criteria proposed in the literature [103,106,107]. The gap between the discourse of co-management and its practical implementation highlights the persistence of top-down control and raises concerns about the timeliness and effectiveness of these mechanisms, especially given the precarious state of clam resources. Moreover, even if these committees were to function as genuine co-management bodies, all decisions and management plans would still require approval from the Directorate General for Maritime Fisheries and the European Commission under the Common Fisheries Policy (CFP; Reg. (EU) No 1380/2013, including the regionalisation provisions of Article 18). The CFP remains the central mechanism of EU fisheries governance, providing a common regulatory framework for member states’ fisheries[108]. Consequently, the overall governance structure remains fundamentally top-down. This underscores an inherent tension between scientific authority and democratic legitimacy, as fisheries management often relies on technocratic, expert-driven decision-making (Linke and Jentoft, 2016; Linke and Siegrist, 2024; Wilson, 2009). In the case of clam fisheries, continued reliance on top-down decision-making allows little flexibility in responding to resource collapse. Unless co-management structures become genuinely binding and meaningfully incorporate fishers’ knowledge into decision-making, management will continue to lag ecological decline. An urgent transformation is therefore needed to restore clam stocks and safeguard the livelihoods of coastal communities. As highlighted by [112], there is growing concern that top-down governance in European SSF delivers “too little, too late”. This issue is particularly pressing in the Northwestern Mediterranean, where delayed regulatory action coincides with multiple stressors, including the accelerating effects of climate change, further jeopardising recovery prospects. Within this broader governance context, the available evidence suggests that EU-driven regulatory frameworks introduced during the 2010s have had uneven effects across regions, largely depending on the capacity of regional administrations to implement effective monitoring and enforcement.

Overall, the case of Spanish clam fisheries highlights the critical interplay between ecological dynamics, regulatory design, and governance modes. Our results suggest that the interaction between ecological conditions and governance frameworks plays a key role in determining whether fisheries recover after periods of decline. In more productive areas, such as the Atlantic coast of Andalusia and parts of the Alboran Sea, clam populations have shown repeated recovery following periods of decline. This resilience may reflect the combination of favourable environmental conditions and the gradual implementation of adaptive management measures. In contrast, less productive regions—particularly the Northwestern Mediterranean—have experienced recurrent collapses and prolonged closures under top-down governance, limited regulatory consistency, and the absence of supra-regional coordination. These findings suggest that sustaining small-scale clam fisheries requires a shift away from reactive management and fragmented jurisdictional strategies towards coordinated supra-regional frameworks that align with biological population structures and integrate scientific assessments with fishers’ ecological knowledge. This transformation is particularly urgent in the Northwestern Mediterranean, where overexploitation, low productivity, governance limitations, and climate change interact to create a highly complex problem. Continued attention is nevertheless required even in the more productive regions to ensure that SSF remain resilient and that present management practices do not erode long-term sustainability.

## 5. Acknowledgments

We are grateful to the Departament d’Agricultura, Ramaderia i Pesca de la Generalitat de Catalunya; to Dr Jose Valencia from the Balearic Islands; to the Conselleria d’Agricultura, Aigua, Ramaderia i Pesca de la Generalitat Valenciana; to the Consejería de Agua, Agricultura, Ganadería y Pesca de la Región de Murcia; and to the Consejería de Agricultura, Pesca, Agua y Desarrollo Rural (Junta de Andalucía) for providing the fisheries data. We also thank the anonymous reviewers for their constructive comments, which helped improve this manuscript.

## 6. Funding sources

This work was supported by the Programa de Ayudas a Proyectos de I+D+I del Plan Complementario de Ciencias Marinas y del Plan de Recuperación, Transformación y Resiliencia de la Comunidad Autónoma de Andalucía (Project PCM_00122 to CR and MH). This study forms part of the ThinkInAzul programme supported by MCIN with funding from the European Union Next Generation EU (PCM_00122 and PRTR-C17.I1).

## 7. Declaration of Interest statement

The authors declare that they have no known competing financial interests or personal relationships that could have appeared to influence the work reported in this manuscript.

## 8. Data Availability

Data will be made available on request.

## Supplementary Material: Key legislation on clam fisheries (1980s-2020s) by CCAA

### Catalonia

- **Order 20/09/1985**. Established minimum legal size (MLS) for *Donax trunculus* (27.2 mm length, 8.3 mm width).
- **Decree 9/1987**. Defined user groups authorised for mechanised clam dredges (winch-operated) and hand-operated dredges; created separate licensing frameworks.
- **Order 21/07/1987 (Roses)**. Introduced specific appropriation rules for Roses area (access, zoning, depth, quotas, vessel specifications).
- **Order 28/01/1988 (North Barcelona)**. Detailed regulations for fishing grounds in North Barcelona.
- **Orders 09/06/1993; 10/05/1994; 20/09/2000; AAM/89/2011.** Defining production areas for clams along the coast.
- **Order 13/06/1994 (Ebro Delta)**. Appropriation rules to resolve conflicts between mechanised dredges and hand gatherers.
- Orders 01/07/1987; 26/10/1994; 31/07/1995; 03/10/1997; 10/08/1999; 18/10/1999. Technical specifications for vessel-towed dredges (*Bolinus brandaris* as main target, up to 10% bycatches including clams).
- **Order 24/12/2001**. Increased allowable bycatches for vessel-towed dredges from 10% to 20%.
- **Order ARP/92/2002 (Roses)**. Temporary biological closure and restricted hours (repealed by **Order ARP/169/2003**).
- **Order AAR/143/2007**. Unified technical standards for vessels and mechanised clam dredges across Catalonia (mesh size, vessel specs).
- **Order AAR/60/2008 (North Barcelona)**. Temporary closure and reduced daily fishing hours for mechanised dredges.
- **Orders AAM/127/2014; ARP/256/2016; ARP/182/2019**. Management plans for hand-operated dredges.
- **Orders ARP/362/2015; ARP/122/2020.** Management plans for mechanised clam dredges.
- **Orders ARP/114/2018; ARP/188/2020**. Management plans for vessel-towed dredges.

### Valencian Community

- **Order 15/09/1988**. Introduced a clam fishing license; formally distinguished between vessel-based mechanised dredges and hand-operated dredges.
- **Resolution 02/06/1993**. Defining production areas for clams along the coast.
- **Order 07/06/1993**. Provided the initial legal framework for regional management; set limits on dredge mouth width (75 cm mechanised, 85 cm hand-operated), number of dredges per vessel (up to four), onboard sieving mesh sizes (19 mm for *Chamelea gallina*, 14 mm for *D. trunculus*), and daily catch limits; restricted the number of hand-gathering permits per fishermen’s guild (cofradía) to 25.
- **Resolutions 20/01/1995;12/02/1996**; and **13/04/2007**. Establishing working hours for shellfishing activity along the Valencian coast.
- **Decree 67/1996**. Introduced an official clam fishing license, consolidating the distinction between mechanised dredges and hand-operated dredges, thereby creating separate user groups.
- **Decree 94/2013 (12 July)**. Updated the legal framework for clam fisheries; defined user eligibility, fishing schedules, technical gear specifications, minimum legal sizes, and daily TACs per vessel or shell fisher.
- **Resolution 09/08/2013**. Imposed indefinite closures for *C. gallina* and *D. trunculus* in three fishing grounds: Patagona-Cabanyal, Molinell-Devenes, and Tavernes.
- **Decree 62/2016 (20 May).** Established the first comprehensive clam fishery Management Plan; included rules for mechanised and hand-operated dredges, TACs, MLS, daily catch limits, and schedules.
- **Resolution 18/05/2017**. Extended the indefinite closure established in 2013 to cover the entire Valencian coast.

### Balearic Islands

- **Order 24/02/1993** – Formally regulated the clam fishery; introduced a mandatory licensing system; set daily catch limits based on crew size (15-20 kg for one fisher; 40-60 kg for three); established a TAC of 70 tonnes; restricted vessels to two dredges; capped daily working hours at nine (later reduced to seven); and specified technical dredge requirements and onboard mesh sizes.
- **Order 31/10/1994** – Updated appropriation rules, including fishing schedules and effort controls.
- **Resolution 10/11/1995**. Annual update to appropriation rules.
- **Resolution 21/10/1996**. Annual update to appropriation rules.
- **Resolution 10/10/1997**. Annual update to appropriation rules.
- **Resolution 16/10/1998**. Annual update to appropriation rules.
- **Order 7/02/2001.** Establishing mollusc and other invertebrate production zones in the Balearic Islands.
- **Order 20/12/2002.** Regional government implemented precautionary closure of the Palma Bay clam fishery in response to declining landings.
- **Orders 07/02/2001; 28/03/2003**. Defining production areas for clams along the coast.
- **Resolution 28/08/2003.** Approving the call for aid for the temporary cessation of the wedge clam (*C. gallina*) fishery in Palma Bay.

### Murcia

- **Resolution 09/04/1984**. Introduced minimum legal size (MLS) of 18 mm and temporary closure (April-July 1985).
- **Resolution 10/04/1985**. Lifted previous closure.
- **Order 09/04/1986**. Reinstated closure (April 1986-May 1987); raised MLS to 20 mm; regulated users, dredge specifications, and fishing schedules (07:00-13:00).
- **Order 09/09/1987**. Extended fishing period by one month.
- **Order 12/05/1988**. Shortened fishing season to June only; reduced fishing hours (07:00-11:00); imposed TACs of 20 kg per crew member/day.
- **Order 10/04/1990**. Expanded season (15 April-15 May); extended fishing hours.
- **Order 28/05/1991**. Lengthened fishing season by one additional month.
- **Order 12/06/1992**. Reduced fishing season (15 April-20 June); restricted hours (08:00-11:00); raised MLS to 21 mm; limited dredges per vessel; mandated sorting at guild markets.
- **Resolution 13/07/1994**. Repealed closure of Cabos Negrete-Palos before enforcement.
- **Resolution 30/06/1995**. Indefinitely closed Cabos Negrete-Palos fishing ground.
- **Order 16/06/2002**. Definitively closed all clam fishing grounds in Murcia; fishery permanently shut down.

### Andalusia

- **Order 16/02/1984.** MLS: 24 mm (*C. gallina*), 30 mm (*D. trunculus*), 60 mm (*C. chione*); seasonal closures (6-8 months) (all Andalusia).
- **Order 19/11/1984.** Specific shell-fishing license introduced (all Andalusia).
- **Resolution 01/03/1985.** Adjusted closures; daily catch limits (150 kg/vessel) (Mediterranean, Algeciras-Punta Chullera).
- **Order 23/11/1986**. Census of the shellfishing fleet (all Andalusia).
- **Order 25/04/1993.** Delineation of clam fishing areas (all Andalusia).
- **Order 24/10/1994.** Update of shellfishing fleet census (all Andalusia).
- **Order 24/06/1996.** Regulated towed dredges for *C. gallina* (technical specs, 300 kg/day, MLS 24 mm) (Atlantic).
- **Order 22/12/1998.** Defined geographic boundaries of fishing grounds (all Andalusia).
- **Order 25/03/1999**. Regulated hydraulic dredges for *C. gallina* (MLS 26 mm, depth > 5 m, seasonal closure) (Atlantic).
- **Order 09/11/1999.** Updated delineation of clam fishing areas (all Andalusia).
- Order 28/01/2000. Regulation of striped venus clam (*C. gallina*) fishery in the Gulf of Cádiz (Atlantic).
- **Order 25/03/2003.** Raised MLS for *C. gallina* to 25 mm; minor closure updates (all Andalusia).
- **Order 16/06/2004**. Established Guadalquivir estuary reserve; zoning for dredges and hand gathering (Atlantic).
- **Order 11/01/2005.** Specific rules for subzones in the Guadalquivir reserve (Atlantic).
- **Resolution 02/05/2007**. List of vessels authorised in Guadalquivir Reserve Zones B and C (Atlantic).
- **Order 24/09/2008**. Regulation of professional on-foot shellfishing licenses (all Andalusia).
- **Order 23/09/2008.** Census of shellfishing vessels targeting bivalves and gastropods (all Andalusia).
- **Order 22/04/2010.** Modified on-foot shellfishing licensing (all Andalusia).
- **Decree 387/2010.** Distinguished on-foot vs vessel-based shellfishing; licensing schemes (all Andalusia).
- **Order 06/07/2010**. Modified Guadalquivir estuary reserve rules (Atlantic).
- **Order 16/12/2010.** Precautionary closure of Guadalquivir *C. gallina* fishery (Atlantic).
- **Order 01/04/2011**. Created shellfishing reserve in Huelva; modified on-foot shellfishing licenses (all Andalusia).
- **Order 24/06/2011.** Regulating shellfishing from vessels using hydraulic dredges in the Gulf of Cádiz and establishing a fishing effort adjustment plan for the fleet operating under this modality and fishing ground.
- **Order 28/09/2011.** Regulated vessel-based shellfishing with towed dredges in the Gulf of Cádiz.
- **Decree 64/2012.** Regulated working days, hours and vessel monitoring (all Andalusia).
- **Order 26/02/2013.** Standardised working hours across modalities (all Andalusia).
- **Order 13/06/2013.** Regulated fishing schedules (Atlantic).
- **Order 24/03/2014.** Management plan for mechanised dredges (Mediterranean).
- **Decree 99/2015.** Reinforced separation on-foot/vessel-based; licensing (all Andalusia).
- **Order 30/11/2016.** Temporary closure of *C. gallina* in Guadalquivir (Atlantic).
- **Order 29/06/2017.** Comprehensive regulatory framework for *C. gallina*; VMS, quotas, monitoring (Atlantic).
- **Order 23/11/2017.** Standardised working hours (all Andalusia).
- **Order 22/02/2018.** Reduced MLS for *D. trunculus* to 25 mm (all Andalusia).
- **Order 27/04/2018**. Adaptation of production zones of bivalves and invertebrates; controls (all Andalusia).
- **Resolution 24/09/2018.** Opening of *C. gallina* fishery in Gulf of Cádiz (Atlantic).
- **Resolution 01/10/2018.** Partial modification of Resolution 24/09/2018 (Atlantic).
- **Resolution 09/01/2019.** Sanitary classification of production zones (all Andalusia).
- **Resolution 30/01/2019.** Partial modification of Resolution 24/09/2018 (Atlantic).
- **Order 27/03/2019.** Modified hydraulic dredges regulation (Atlantic).
- **Resolution 04/04/2019.** Closure of *C. gallina* fishery in Gulf of Cádiz (Atlantic).
- **Resolution 21/06/2019.** Fishing plan for *C. gallina* 2019-2020 (Atlantic).
- **Order 27/12/2019.** Management plan for mechanised dredges (Mediterranean).
- **Resolution 11/03/2020.** Closure of *C. gallina* fishery (Atlantic).
- **Order 06/04/2020.** Dedicated management plan for *C. gallina* (Atlantic).
- **Resolution 03/04/2020.** Adjusted closure for *D. trunculus* (all Andalusia).
- **Resolution 25/06/2020.** Fishing plan for *C. gallina* 2020-2021 (Atlantic).
- **Resolution 22/07/2020.** Modified fishing plan for *C. gallina* 2020-2021 (Atlantic).
- **Resolution 15/09/2020**. Closure of *C. gallina* in production zone (Atlantic).
- **Order 25/05/2020**. Regulatory framework for on-foot *D. trunculus* (Atlantic).
- **Resolution 19/03/2021.** Sanitary classification of production zones (all Andalusia).
- **Order 28/01/2021**. Call for new on-foot shellfishing licences in Huelva and Cádiz.
- **Resolution 08/02/2021**. Temporary closure *C. gallina* (Atlantic).
- **Resolution 18/02/2021**. Closure of *C. gallina* fishery (Atlantic).
- **Resolution 23/06/2021.** Fishing plan for *C. gallina* 2021-2022 (Atlantic).
- **Resolution 22/02/2022.** Closure of *C. gallina* fishery (Atlantic).
- **Order 29/06/2022.** Modified Guadalquivir reserve rules (Atlantic).
- **Resolution 28/06/2022.** Fishing plan for *C. gallina* 2022-2023 (Atlantic).
- **Resolution 16/03/2023.** Temporary closure *C. gallina* (Atlantic).
- **Resolution 27/04/2023.** Call for fleet data submission (Atlantic).
- **Resolution 13/07/2023.** Fishing plan for *C. gallina* 2023–2024; extended 2020 plan (Atlantic).
- **Resolution 31/07/2023.** Opening *C. gallina* in zone AND-102 Barra del Terrón; modified technical measures (Atlantic).
- **Resolution 25/08/2023.** Opening *C. gallina* in zone AND-107 Doñana Norte; technical adjustments (Atlantic).

---

**Figure S1.**
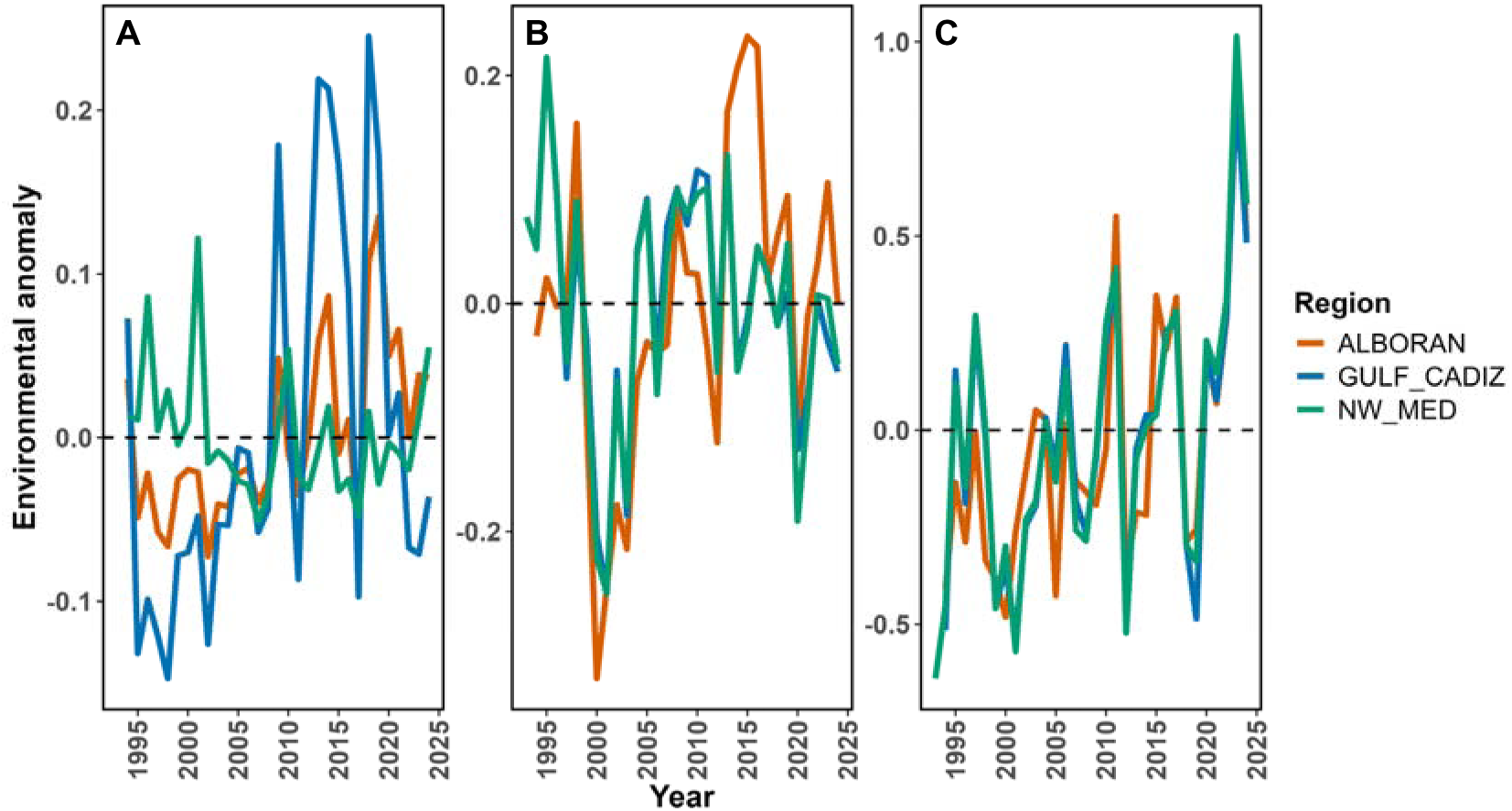
Annual anomalies of **(A)** sea surface temperature (SST), **(B)** sea surface salinity (SSS), and **(C)** chlorophyll-a (Chl) for the three study regions (NW Mediterranean, Alboran Sea, and Gulf of Cadiz) during 1993-2024. Environmental data was obtained from Copernicus Marine Service products and aggregated annually. Anomalies represent deviations from the long-term mean of the study period.

**Table S1.** Pearson correlation coefficients (r) and associated p-values between annual landings and environmental variables (SST lagged one year, SSS lagged one year, and chlorophyll-a anomaly) for each species–region combination. Significant correlations (p < 0.05) are shown in bold. Environmental data were obtained from Copernicus Marine Service datasets and aggregated annually.

| Species | Region | $r_{\text{SST}}$ | $p_{\text{SST}}$ | $r_{\text{SSS}}$ | $p_{\text{SSS}}$ | $r_{\text{CHL}}$ | $p_{\text{CHL}}$ |
| --- | --- | --- | --- | --- | --- | --- | --- |
| *Chamelea gallina* | ALBORAN | -0.33 | 0.076 | -0.05 | 0.794 | -0.05 | 0.771 |
| *Chamelea gallina* | GULF_CADIZ | -0.34 | 0.070 | -0.25 | 0.188 | 0.26 | 0.155 |
| *Chamelea gallina* | NW_MED | **-0.37** | **0.043** | 0.29 | 0.117 | 0.35 | 0.055 |
| *Donax trunculus* | ALBORAN | -0.04 | 0.854 | 0.02 | 0.910 | 0.17 | 0.349 |
| *Donax trunculus* | GULF_CADIZ | 0.23 | 0.222 | -0.20 | 0.295 | 0.11 | 0.571 |
| *Donax trunculus* | NW_MED | **-0.42** | **0.020** | 0.16 | 0.390 | 0.20 | 0.277 |
| *Callista chione* | ALBORAN | **-0.45** | **0.013** | -0.18 | 0.333 | -0.12 | 0.514 |
| *Callista chione* | NW_MED | -0.20 | 0.287 | 0.33 | 0.069 | **0.45** | **0.012** |

## Notes

### Competing Interest Statement

The authors have declared no competing interest.

### Summary of Updates

The revised manuscript addresses the comments of the referees raised in the first round of peer review in Marine Policy by strengthening both structure and analysis. It now presents a clearer comparative socio-ecological framework, expands the description of governance methods, and rewrites the Introduction to better frame the study around interactions between ecological drivers and fisheries management. The breakpoint analysis was removed, and an environmental analysis was added to examine links between landings and sea surface temperature, salinity, and chlorophyll-a. The manuscript also clarifies that the NW Mediterranean grouping is only for statistical purposes, not a claim of shared governance. The Discussion was reorganised to foreground the main findings, broaden regional comparisons, and better distinguish the roles of environmental change and governance. Figure legends and interpretive passages were also clarified, improving readability and overall coherence.

## Bibliography

[1] FAO, The State of World Fisheries and Aquaculture 2024 – Blue Transformation in action., Rome, 2024.

[2] C.Y. Chan, N. Tran, Y. Hoong, T.B. Sulser, Y.M. Aung, What do we know about the future of aquatic foods in global agri-food systems? Initiative on Resilient Aquatic Foods Systems., 2023.

[3] Eurostat, Fisheries statistics: Aquaculture production and fishery imports in the EU., (2025).

[4] UMOFA, The EU fish market, 2024.

[5] J. Lloret, I.G. Cowx, H. Cabral, M. Castro, T. Font, J.M.S. Gonçalves, A. Gordoa, E. Hoefnagel, S. Matić-Skoko, E. Mikkelsen, B. Morales-Nin, D.K. Moutopoulos, M. Muñoz, M.N. dos Santos, P. Pintassilgo, C. Pita, K.I. Stergiou, V. Ünal, P. Veiga, K. Erzini, Small-scale coastal fisheries in European Seas are not what they were: Ecological, social and economic changes, Mar. Policy 98 (2018) 176–186. 10.1016/j.marpol.2016.11.007.

[6] M. Baeta, M.A. Solís, M. Ballesteros, O. Defeo, Long-term trends in striped venus clam (Chamelea gallina) fisheries in the western Mediterranean Sea: The case of Ebro Delta (NE Spain), Mar. Policy 134 (2021). 10.1016/j.marpol.2021.104798.

[7] M. Baeta, M. Ramón, E. Galimany, Decline of a Callista chione (Bivalvia: Veneridae) bed in the Maresme coast (northwestern Mediterranean Sea), Ocean Coast. Manag. 93 (2014) 15–25. 10.1016/j.ocecoaman.2014.03.001.

[8] M. Baeta, M.A. Solís, S. Frias-Vidal, L. Claramonte, A. Sepouna, M. Ballesteros, The wedge clam (Donax trunculus) hand-operated fishery in the NW Mediterranean Sea: Landings, catch composition, damage rates and impact of fishing activity, Ocean Coast. Manag. 237 (2023) 106534. 10.1016/j.ocecoaman.2023.106534.

[9] D. Escobar-Ortega, N. Fernández, L. Couceiro, R. Muíño, P. Pita, E. Martínez, D. Fernández-Márquez, Polychaete exploitation in Galicia (NW Spain): Challenges, advances, and pathways for sustainable resource management, Ocean Coast. Manag. 256 (2024) 107302. 10.1016/J.OCECOAMAN.2024.107302.

[10] B. Lopez, Marginal marine crustacean fisheries, Fisheries and Aquaculture. Oxford University Press, Oxford (2020) 159–180.

[11] M. Baeta, M.A. Solís, S. Frias-Vidal, L. Claramonte, A. Sepouna, M. Ballesteros, The wedge clam (Donax trunculus) hand-operated fishery in the NW Mediterranean Sea: Landings, catch composition, damage rates and impact of fishing activity, Ocean Coast. Manag. 237 (2023). 10.1016/j.ocecoaman.2023.106534.

[12] B. Crawford, M.D. Herrera, N. Hernandez, C.R. Leclair, N. Jiddawi, S. Masumbuko, M. Haws, Small Scale Fisheries Management: Lessons from Cockle Harvesters in Nicaragua and Tanzania, Coastal Management 38 (2010) 195–215. 10.1080/08920753.2010.483174.

[13] O. Guyader, P. Berthou, C. Koutsikopoulos, F. Alban, S. Demanèche, M.B. Gaspar, R. Eschbaum, E. Fahy, O. Tully, L. Reynal, O. Curtil, K. Frangoudes, F. Maynou, Small scale fisheries in Europe: A comparative analysis based on a selection of case studies, Fish. Res. 140 (2013) 1–13. 10.1016/J.FISHRES.2012.11.008.

[14] J. Pascual-Fernandez, C. Pita, M. Bavinck, Small-Scale Fisheries in Europe: Status, Resilience and Governance, 2020. 10.1007/978-3-030-37371-9.

[15] J.L. Jacquet, D. Pauly, The rise of seafood awareness campaigns in an era of collapsing fisheries, Mar. Policy 31 (2007) 308–313. 10.1016/j.marpol.2006.09.003.

[16] C. Pita, J.J. Pascual-Fernández, M. Bavinck, Small-scale fisheries in Europe: challenges and opportunities, Small-Scale Fisheries in Europe: Status, Resilience and Governance (2020) 581–600.

[17] F. Natale, N. Carvalho, A. Paulrud, Defining small-scale fisheries in the EU on the basis of their operational range of activity The Swedish fleet as a case study, Fish. Res. 164 (2015) 286–292. 10.1016/j.fishres.2014.12.013.

[18] M. Delgado, L. Silva, S. Román, S. Llorens, A. Rodríguez-Rúa, M. Cojan, M. Hidalgo, Spatial distribution patterns of striped venus clam (Chamelea gallina, L. 1758) natural beds in the Gulf of Cádiz (SW Spain): Influence of environmental variables and management considerations, Reg. Stud. Mar. Sci. 63 (2023) 103024. 10.1016/j.rsma.2023.103024.

[19] M.P. Sastre, Aggregated patterns of dispersion in Donax denticulatus, Bull. Mar. Sci. 36 (1985) 220–224.

[20] O. Defeo, A. McLachlan, Patterns, processes and regulatory mechanisms in sandy beach macrofauna: a multi-scale analysis, Mar. Ecol. Prog. Ser. 295 (2005) 1–20.

[21] M. Delgado, L. Silva, S. Gómez, E. Masferrer, M. Cojan, M.B. Gaspar, Population and production parameters of the wedge clam Donax trunculus (Linnaeus, 1758) in intertidal areas on the southwest Spanish coast: Considerations in relation to protected areas, Fish. Res. 193 (2017) 232– 241. 10.1016/j.fishres.2017.04.012.

[22] J.M. Pérès, J. Picard, Nouveau manuel de bionomie benthique de la Mer Méditerranée, Recueil Des Travaux de La Station Marine d’Endoume (1964) 31–47.

[23] R. Carlucci, G. Cipriano, D. Cascione, M. Ingrosso, E. Barbone, N. Ungaro, P. Ricci, Influence of hydraulic clam dredging and seasonal environmental changes on macro-benthic communities in the Southern Adriatic (Central Mediterranean Sea), BMC Ecol. Evol. 24 (2024). 10.1186/S12862-023-02197-9.

[24] C. García-Fernández, C. Ciércoles, J. Urra, P. Marina, J.M. Serna-Quintero, J. Baro, Influence of environmental conditions on the abundance of the striped venus Chamelea gallina in the northern Alboran Sea (Western Mediterranean Sea), Reg. Stud. Mar. Sci. 76 (2024) 103601. 10.1016/j.rsma.2024.103601.

[25] D. Burdon, R. Callaway, M. Elliott, T. Smith, A. Wither, Mass mortalities in bivalve populations: A review of the edible cockle Cerastoderma edule (L.), Estuar. Coast. Shelf Sci. 150 (2014) 271–280. 10.1016/j.ecss.2014.04.011.

[26] S.M. Fiori, N.J. Cazzaniga, Mass mortality of the yellow clam, Mesodesma mactroides (Bivalvia: Mactracea) in Monte Hermoso beach, Argentina, Biol. Conserv. 89 (1999) 305–309. 10.1016/S0006-3207(98)00151-7.

[27] H.F.E. Jones, C.A. Pilditch, D.P. Hamilton, K.R. Bryan, Impacts of a bivalve mass mortality event on an estuarine food web and bivalve grazing pressure, N. Z. J. Mar. Freshwater Res. 51 (2017) 370–392. 10.1080/00288330.2016.1245200.

[28] M.B. Gaspar, A.M. Pereira, P. Vasconcelos, C.C. Monteiro, Age and growth of Chamelea gallina from the Algarve coast (Southern Portugal): influence of seawater temperature and gametogenic cycle on growth rate, Journal of Molluscan Studies 70 (2004) 371–377. 10.1093/mollus/70.4.371.

[29] M.M. Zeichen, S. Agnesi, A. Mariani, A. Maccaroni, G.D. Ardizzone, Biology and Population Dynamics of Donax trunculus L. (Bivalvia: Donacidae) in the South Adriatic Coast (Italy), Estuar. Coast. Shelf Sci. 54 (2002) 971–982. 10.1006/ecss.2001.0868.

[30] M. Baeta, F. Breton, R. Ubach, E. Ariza, A socio-ecological approach to the declining Catalan clam fisheries, Ocean Coast. Manag. 154 (2018) 143–154. 10.1016/j.ocecoaman.2018.01.012.

[31] M. Baeta, M. Solís, M. Ramón, M. Ballesteros, Effects of fishing closure and mechanized clam dredging on a Callista chione bed in the western Mediterranean Sea, Reg. Stud. Mar. Sci. 48 (2021) 102063. 10.1016/J.RSMA.2021.102063.

[32] J. Escrivá, M. Rodilla, F. Llario, S. Falco, The Collapse of a Wedge Clam Fishery in the Spanish Mediterranean Coast and Recovery Problems, J. Shellfish Res. 40 (2021) 37–47. 10.2983/035.040.0105.

[33] M.Á. Nombela, P. Diz, E.N. Couto, G. Martínez, Textural Characteristics might Influence Donax trunculus Shellfishing Banks Exploitability, Thalassas: An International Journal of Marine Sciences 33 (2017) 87–93. 10.1007/s41208-017-0025-2.

[34] M. Fabinyi, K. Barclay, Fisheries Governance, in: 2021: pp. 65–90. 10.1007/978-3-030-79591-7_4.

[35] A.D. Marie, C. Lejeusne, E. Karapatsiou, J.A. Cuesta, P. Drake, E. Macpherson, L. Bernatchez, C. Rico, Implications for management and conservation of the population genetic structure of the wedge clam Donax trunculus across two biogeographic boundaries, Sci. Rep. 6 (2016) 39152. 10.1038/srep39152.

[36] J. Escrivá Perales, Estudio de los bancos explotables de donax trunculus y chamelea gallina en el sector sur del golfo de valencia y factores ambientales que influyen en su abundancia, Universitat Politècnica de València, 2019.

[37] J. Urra, T. García, H. Gallardo-Roldán, E. León, M. Lozano, J. Baro, J.L. Rueda, Discard analysis and damage assessment in the wedge clam mechanized dredging fisheries of the northern Alboran Sea (W Mediterranean Sea), Fish. Res. 187 (2017) 58–67. 10.1016/j.fishres.2016.10.018.

[38] M. Ramón, Estudio de la poblaciones de Chamelea gallina (Linnaeus, 1758) y Donax trunculus (Linnaeus, 1758) (MolluscaL:Bivalva) en el Golfo de Valencia (Mediterráneo Occidental), Universitat de Barcelona, 1993.

[39] Solytur, Monitor de competividad turística responsable de los destinos de “sol y playa” español, 2024. https://www.exceltur.org/wp-content/uploads/2025/04/Exceltur_Solytur-2024_230425.pdf (accessed October 10, 2025).

[40] UNEP/MAP, State of the Mediterranean. Marine and Coastal Environment, UNEP/MAP – Barcelona Convention. Athens, 2012. 10.13140/RG.2.1.3013.2648.

[41] A. Penaud, F. Eynaud, J. Etourneau, J. Bonnin, A. de Vernal, S. Zaragosi, J.-H. Kim, S. Kang, J.- K. Gal, D. Oliveira, C. Waelbroeck, Ocean Productivity in the Gulf of Cadiz Over the Last 50 kyr, Paleoceanogr. Paleoclimatol. 37 (2022) e2021PA004316. 10.1029/2021PA004316.

[42] L. Prieto, G. Navarro, S. Rodríguez-Gálvez, I.E. Huertas, J.M. Naranjo, J. Ruiz, Oceanographic and meteorological forcing of the pelagic ecosystem on the Gulf of Cadiz shelf (SW Iberian Peninsula), Cont. Shelf Res. 29 (2009) 2122–2137. 10.1016/j.csr.2009.08.007.

[43] A. García de Vinuesa, D. Florido, C. Vilas, M.Á. Torres, M. Delgado, I. Muñoz, R. Cabrera-Castro, F. Ramos, M. Llope, Framing social systems for ecosystem-based management: The Guadalquivir estuary-Gulf of Cadiz coupled SES as case study, Environ. Dev. 55 (2025) 101206. 10.1016/j.envdev.2025.101206.

[44] A. Peris, Y. Soriano, Y. Picó, M.A. Bravo, G. Blanco, E. Eljarrat, Pesticides in water and sediments from natural protected areas of Spain and their associated ecological risk, Chemosphere 362 (2024) 142628. 10.1016/j.chemosphere.2024.142628.

[45] I. Caballero, E.P. Morris, L. Pietro, G. Navarro, The influence of the Guadalquivir river on spatio-temporal variability in the pelagic ecosystem of the eastern Gulf of Cádiz, (2014).

[46] M. Llope, The ecosystem approach in the Gulf of Cadiz. A perspective from the southernmost European Atlantic regional sea, ICES Journal of Marine Science 74 (2017) 382–390. 10.1093/icesjms/fsw165.

[47] T. Oguz, D. Macías Moy, J. García-Lafuente, P. Ananda, J. Tintoré, Fueling Plankton Production By a Meandering Frontal Jet: A Case Study For The Alboran Sea (Western Mediterranean), PLoS One 9 (2014) e111482. 10.1371/journal.pone.0111482.

[48] R. Carlucci, D. Cascione, P. Ricci, D. De Padova, V. Dragone, G. Cipriano, M. Mossa, Fluctuations in abundance of the striped venus clam Chamelea gallina in the southern Adriatic Sea (Central Mediterranean Sea): knowledge, gaps and insights for ecosystem-based fishery management, Rev. Fish Biol. Fish 34 (2024) 827–848. 10.1007/S11160-024-09840-8.

[49] M. Dağtekin, C.E. Özyurt, About striped venus clam (Chamelea gallina) fisheries in the Black Sea: Management and economic aspects, Reg. Stud. Mar. Sci. 60 (2023) 102819. 10.1016/j.rsma.2023.102819.

[50] S. Joo, K. Jo, H. Bae, H. Seo, T. Kim, Optimal sediment grain size and sorting for survival and growth of juvenile Manila clams, Venerupis philippinarum, Aquaculture 543 (2021) 737010. 10.1016/j.aquaculture.2021.737010.

[51] S. Joaquim, M.B. Gaspar, D. Matias, R. Ben-Hamadou, W.S. Arnold, Rebuilding viable spawner patches of the overfished Spisula solida (Mollusca: Bivalvia): a preliminary contribution to fishery sustainability, ICES Journal of Marine Science 65 (2008) 60–64. 10.1093/icesjms/fsm167.

[52] O. Defeo, A. de Alava, Effects of human activities on long-term trends in sandy beach populations: the wedge clam Donax hanleyanus in Uruguay, Mar. Ecol. Prog. Ser. 123 (1995) 73–82. https://www.int-res.com/abstracts/meps/v123/meps123073.

[53] C. Goutelle, Tellines: disparition énigmatique d’un coquillage emblématique, La Brèche (2023).

[54] D. Biotope P2A, Étude globale sur la telline en Camargue - Parc Naturel Régional de Camargue Donax trunculus (Linné 1767), 2007.

[55] D. Martínez-Patiño, S. Nóvoa, J. Ojea, M.Á. Nombela, I. Alejo, M.J. Álvarez-Fernández, X.A. Álvarez-Salgado, J. Oetro, O. Teijeiro, J.D. Cerdeira-Arias, I.M. Liñán, C. Brezmes, F. Febrero, L.M.R. González, Determinación de las causas de la disminución de los bancos de coquina (Donax sp.) en Galicia. Banco de Vilarrube, (2019).

[56] M. Ben-Haddad, S. Hajji, M.R. Abelouah, A. Ait Alla, Urbanization Compromises the Sustainability of Coastal Ecosystems: Insights from the Reproductive Traits of the Bioindicator Clam Donax trunculus, Sustainability 17 (2025). 10.3390/su17146622.

[57] G. Bargione, F. Donato, S. o Guizzardi, M.J.M. Macchia, D. Li Veli, A. Lucchetti, Integrating biological traits of Ensis minor into regional management plans: a case for adaptive fisheries policies, Front. Mar. Sci. Volume 12–2025 (2025). 10.3389/fmars.2025.1717257.

[58] M. Romanelli, C.A. Cordisco, O. Giovanardi, The long-term decline of the Chamelea gallina L.(Bivalvia: Veneridae) clam fishery in the Adriatic Sea: is a synthesis possible?, Acta Adriat. 50 (2009) 171–205.

[59] H. Bouzaidi, O. Haroufi, A. Khaili, C. El Fanichi, M. Maatouk, B. El-Moumni, M. Daoudi, Stock assessment and spatial distribution of the smooth clam Callista chione (Linnaeus, 1758) exploited in the occidental Mediterranean Sea of Morocco, Aquaculture, Aquarium, Conservation & Legislation 13 (2020) 1268–1284.

[60] M.J. Fogarty, L.W. Botsford, Population connectivity and spatial management of marine fisheries, Oceanography 20 (2007) 112–123.

[61] C. Strasser, Metapopulation dynamics of the softshell clam, Mya arenaria, (2008). 10.1575/1912/2323.

[62] S.D. Gaines, B. Gaylor, L.R. Gerber, A. Hastings, B.P. Kinlan, Connecting places: the ecological consequences of dispersal in the sea, Oceanography 20 (2007) 90–99.

[63] A.L. Shanks, Pelagic Larval Duration and Dispersal Distance Revisited, Biol. Bull. 216 (2009) 373–385. 10.1086/BBLv216n3p373.

[64] C. Froglia, Observations on the growth of Chamelea gallina (L.) and Ensis minor (Chenu) in the middle Adriatic, Quad Di Lab Di Tecnol Della Pesca 2 (1975) 37–48.

[65] E. Grazioli, C. Guerranti, P. Pastorino, G. Esposito, E. Bianco, E. Simonetti, S. Rainis, M. Renzi, A. Terlizzi, Review of the Scientific Literature on Biology, Ecology, and Aspects Related to the Fishing Sector of the Striped Venus (Chamelea gallina) in Northern Adriatic Sea, J. Mar. Sci. Eng. 10 (2022).

[66] L. Neuberger-Cywiak, Y. Achituv, L. Mizrahi, The ecology of Donax trunculus Linnaeus and Donax semistriatus Poli from the Mediterranean coast of Israel, J. Exp. Mar. Biol. Ecol. 134 (1989) 203–220. 10.1016/0022-0981(89)90070-1.

[67] J.D.M.I.G.J. Pérez-Larruscain, Crecimiento y desarrollo larvario del almejón de sangre Callista chione (Mollusca: Bivalvia) en condiciones de laboratorio., Cienc. Mar. 37 (2011) 271–277.

[68] C. Johnstone, M.P. Rodríguez, Conectividad genética en especies marinas a través del estrecho de Gibraltar y el mar de Alborán, (2023).

[69] J. Tintore, P.E. La Violette, I. Blade, A. Cruzado, A Study of an Intense Density Front in the Eastern Alboran Sea: The Almeria–Oran Front, J. Phys. Oceanogr. 18 (1988) 1384–1397. 10.1175/1520-0485(1988)018<1384:ASOAID>2.0.CO;2.

[70] M. Astraldi, S. Balopoulos, J. Candela, J. Font, M. Gacic, G.P. Gasparini, B. Manca, A. Theocharis, J. Tintoré, The role of straits and channels in understanding the characteristics of Mediterranean circulation, Prog. Oceanogr. 44 (1999) 65–108.

[71] N. Staudenmayer, M. Tyre, L. Perlow, Time to Change: Temporal Shifts as Enablers of Organizational Change, Organization Science 13 (2002) 583–597. 10.1287/orsc.13.5.583.7813.

[72] C. Pedroza, S. Salas, Responses of the fishing sector to transitional constraints: From reactive to proactive change, Yucatan fisheries in Mexico, Mar. Policy 35 (2011) 39–49. 10.1016/j.marpol.2010.08.001.

[73] S.L. Brown, K.M. Eisenhardt, The art of continuous change, Adm. Sci. Q. 42 (1997) 1–34.

[74] E. Edmondson, L. Fanning, Implementing adaptive management within a fisheries management context: a systematic literature review revealing gaps, challenges, and ways forward, Sustainability 14 (2022) 7249.

[75] C.S. Holling, Adaptive environmental assessment and management, John Wiley & Sons, 1978.

[76] C.J. Walters, Adaptive management of renewable resources, Macmillan Publishers Ltd, 1986.

[77] G. McDonald, B. Harford, A. Arrivillaga, E.A. Babcock, R. Carcamo, J. Foley, R. Fujita, T. Gedamke, J. Gibson, K. Karr, J. Robinson, J. Wilson, An indicator-based adaptive management framework and its development for data-limited fisheries in Belize, Mar. Policy 76 (2017) 28–37. 10.1016/j.marpol.2016.11.027.

[78] J.R. Nielsen, D. Manh Son, K.-J. Stæhr, H. Hovgård, N.T. Dieu Thuy, K. Ellegaard, F. Riget, D. Van Thi, P. Giang Hai, Adaptive fisheries management in Vietnam: The use of indicators and the introduction of a multi-disciplinary Marine Fisheries Specialist Team to support implementation, Mar. Policy 31 (2007) 143–152. 10.1016/j.marpol.2006.05.013.

[79] UNESCO, Doñana National Park. World Heritage List – Inscription 685, UNESCO World Heritage Centre (1994).

[80] I.G.A.G.E.; S.L.B.J.M.J. Sobrino, Estudio previo para la delimitación de una reserva de pesca en la desembocadura del Guadalquivir, Consejeria Agricultura y Pesca. Junta Andalucía, Sevilla, 2005.

[81] L. Benestan, Population Genomics Applied to Fishery Management and Conservation, in: 2019: pp. 1–23. 10.1007/13836_2019_66.

[82] L. Bernatchez, M. Wellenreuther, C. Araneda, D.T. Ashton, J.M.I. Barth, T.D. Beacham, G.E. Maes, J.T. Martinsohn, K.M. Miller, K.A. Naish, Harnessing the power of genomics to secure the future of seafood, Trends Ecol. Evol. 32 (2017) 665–680.

[83] H. Reiss, G. Hoarau, M. Dickey□Collas, W.J. Wolff, Genetic population structure of marine fish: mismatch between biological and fisheries management units, Fish and Fisheries 10 (2009) 361–395.

[84] G.A. Begg, K.D. Friedland, J.B. Pearce, Stock identification and its role in stock assessment and fisheries management: an overview, Fish. Res. 43 (1999) 1–8. 10.1016/S0165-7836(99)00062-4.

[85] R. Hilborn, Ecosystem-based fisheries management, Mar. Ecol. Prog. Ser. 274 (2004) 275–278.

[86] E. Kenchington, M. Heino, E.E. Nielsen, Managing marine genetic diversity: time for action?, ICES Journal of Marine Science 60 (2003) 1172–1176.

[87] L.A. Kerr, S.X. Cadrin, A.I. Kovach, Consequences of a mismatch between biological and management units on our perception of Atlantic cod off New England, ICES Journal of Marine Science 71 (2014) 1366–1381. 10.1093/icesjms/fsu113.

[88] A. Aguión, E. Ojea, L. García-Flórez, T. Cruz, J.M. Garmendia, D. Davoult, H. Queiroga, A. Rivera, J.L. Acuña-Fernández, G. Macho, Establishing a governance threshold in small-scale fisheries to achieve sustainability, Ambio 51 (2022) 652–665. 10.1007/s13280-021-01606-x.

[89] P. Mejía-Ruíz, R. Perez-Enriquez, J.A. Mares-Mayagoitia, F. Valenzuela-Quiñonez, Population genomics reveals a mismatch between management and biological units in green abalone (Haliotis fulgens), PeerJ 8 (2020) e9722. 10.7717/peerj.9722.

[90] R. Ouréns, I. Naya, J. Freire, Mismatch between biological, exploitation, and governance scales and ineffective management of sea urchin (Paracentrotus lividus) fisheries in Galicia, Mar. Policy 51 (2015) 13–20. 10.1016/j.marpol.2014.07.015.

[91] O. Guyader, P. Berthou, C. Koustikopoulos, F. Alban, S. Demaneche, M. Gaspar, R. Eschbaum, E. Fahy, O. Tully, L. Reynal, Small-scale coastal fisheries in Europe. Final report, (2007).

[92] J. Freire, A. García-Allut, Socioeconomic and biological causes of management failures in European artisanal fisheries: the case of Galicia (NW Spain), Mar. Policy 24 (2000) 375–384. 10.1016/S0308-597X(00)00013-0.

[93] S. Villasante, The management of the blue whiting fishery as complex social-ecological system: The Galician case, Mar. Policy 36 (2012) 1301–1308. 10.1016/j.marpol.2012.02.013.

[94] I. García-Lorenzo, M.Á. Piñeiro-Antelo, S. Villasante, P. Pita, The cofradías’ role within the Fisheries Local Action Groups system: Implications for small-scale fisheries in Galicia (Spain), Sociol. Ruralis 64 (2024) 415–444. 10.1111/soru.12490.

[95] S. Law 3/2001, Ley (Law) 3/2001, de 26 de marzo, de Pesca Marítima del Estado, Boletín Oficial del Estado n° 75, 2001.

[96] J. Kooiman, M. Bavinck, The governance perspective, Fish for Life: Interactive Governance for Fisheries 3 (2005) 11.

[97] European Commission, Green Paper–Reform of the Common Fisheries Policy, 2009.

[98] M.D. Garza-Gil, M.I. Pérez-Pérez, R. Fernández-González, Governance in small-scale fisheries of Galicia (NW Spain): Moving toward co-management?, Ocean Coast. Manag. 184 (2020) 105013. 10.1016/j.ocecoaman.2019.105013.

[99] J.L. Hatchard, T.S. Gray, From RACs to Advisory Councils: Lessons from North Sea discourse for the 2014 reform of the European Common Fisheries Policy, Mar. Policy 47 (2014) 87–93. 10.1016/j.marpol.2014.02.015.

[100] C. Pita, G.J. Pierce, I. Theodossiou, Stakeholders’ participation in the fisheries management decision-making process: Fishers’ perceptions of participation, Mar. Policy 34 (2010) 1093–1102. 10.1016/j.marpol.2010.03.009.

[101] E.M. Finkbeiner, X. Basurto, Re-defining co-management to facilitate small-scale fisheries reform: An illustration from northwest Mexico, Mar. Policy 51 (2015) 433–441. 10.1016/j.marpol.2014.10.010.

[102] J.R. Nielsen, P. Degnbol, K.K. Viswanathan, M. Ahmed, M. Hara, N.M.R. Abdullah, Fisheries co-management—an institutional innovation? Lessons from South East Asia and Southern Africa, Mar. Policy 28 (2004) 151–160. 10.1016/S0308-597X(03)00083-6.

[103] R.S. Pomeroy, F. Berkes, Two to tango: The role of government in fisheries co-management, Mar. Policy 21 (1997) 465–480. 10.1016/S0308-597X(97)00017-1.

[104] A. Rivera, S. Gelcich, L. García-Florez, J.L. Alcázar, J.L. Acuña, Co-management in Europe: Insights from the gooseneck barnacle fishery in Asturias, Spain, Mar. Policy 50 (2014) 300–308. 10.1016/j.marpol.2014.07.011.

[105] S. Gelcich, T. Hughes, P. Olsson, C. Folke, O. Defeo, M. Fernandez, S. Foale, L. Gunderson, C. Rodriguez-Sickert, M. Scheffer, R. Steneck, J. Castilla, Navigating Transformations in Governance of Chilean Marine Coastal Resources, Proc. Natl. Acad. Sci. U. S. A. 107 (2010) 16794–16799. 10.1073/pnas.1012021107.

[106] N.L. Gutiérrez, R. Hilborn, O. Defeo, Leadership, social capital and incentives promote successful fisheries, Nature 470 (2011) 386–389. 10.1038/nature09689.

[107] S. Jentoft, Introduction, in: D.C. Wilson, J.R. Nielsen, P. Degnbol (Eds.), The Fisheries Co-Management Experience: Accomplishments, Challenges and Prospects, Springer Netherlands, Dordrecht, 2003: pp. 1–14. 10.1007/978-94-017-3323-6_1.

[108] E. Penas-Lado, Quo Vadis common fisheries policy?, John Wiley & Sons, 2019.

[109] S. Linke, S. Jentoft, Ideals, realities and paradoxes of stakeholder participation in EU fisheries governance, Environ. Sociol. 2 (2016) 144–154.

[110] S. Linke, N. Siegrist, Aligning top-down and bottom-up modes of governance? How EU Fisheries Local Action Groups support small-scale fisheries and coastal community development in Sweden, Sociol. Ruralis 64 (2024) 490–509. 10.1111/soru.12452.

[111] D.C. Wilson, The paradoxes of transparency: science and the ecosystem approach to fisheries management in Europe, Amsterdam University Press, 2009.

[112] J. Percy, B. O’Riordan, The EU Common Fisheries Policy and Small-Scale Fisheries: A Forgotten Fleet Fighting for Recognition, in: J.J. Pascual-Fernández, C. Pita, M. Bavinck (Eds.), Small-Scale Fisheries in Europe: Status, Resilience and Governance, Springer International Publishing, Cham, 2020: pp. 23–46. 10.1007/978-3-030-37371-9_2.

